# β-catenin has an ancestral role in cell fate specification but not cell adhesion

**DOI:** 10.1101/520957

**Authors:** Miguel Salinas-Saavedra, Athula H. Wikramanayake, Mark Q Martindale

**Affiliations:** The Whitney Laboratory for Marine Bioscience, and the Department of Biology, University of Florida, 9505 N, Ocean Shore Blvd, St. Augustine, FL 32080-8610, USA; Department of Biology, University of Miami, Coral Gables, FL 33124, USA

## Abstract

The ß-catenin protein has two major known functions in animal cells. It keeps epithelial tissue homeostasis by its connection with Adherens Junctions (AJ), and it serves as a transcriptional cofactor along with Lef/Tcf to enter the nucleus and regulate target genes of the Wnt/ß-catenin (cWnt) signaling pathway. To assess the ancestral role of ß-catenin during development we examined its distribution and function in the ctenophore *Mnemiopsis leidyi* (one of the earliest branching animal phyla) by using ctenophore-specific antibodies and mRNA injection. We found that ß-catenin protein never localizes to cell-cell contacts during embryogenesis as it does in other metazoans, most likely because ctenophore-cadherins do not have the cytoplasmic domain required for interaction with the catenin proteins. Downregulation of zygotic *Ml*ß-catenin signaling led to the loss of endodermal and mesodermal tissues indicating that nuclear ß-catenin may have a deep role in germ-layer evolution. Our results indicate that the ancestral role for ß-catenin was in the cell-fate specification and not in cell adhesion and also further emphasizes the critical role of this protein in the evolution of tissue layers in metazoans.

## BACKGROUND

Animal evolutionary history has generated an intriguing diversity of body plans. Changes in developmental programs within and between animal lineages are required to generate this diversity. Within these changes, different modes of gastrulation are observable between major animal groups. For example, the site of gastrulation is specified at the vegetal pole in bilaterians, while in cnidarians and ctenophores it is specified at the animal pole (Figure 1) (*1–3*). Decades of research suggests that this shift of the site of gastrulation in animal evolution may be related to major body plan transitions due to its early role in germ cell-layer specification and embryonic body patterning (*1–4*).

**Figure 1.**
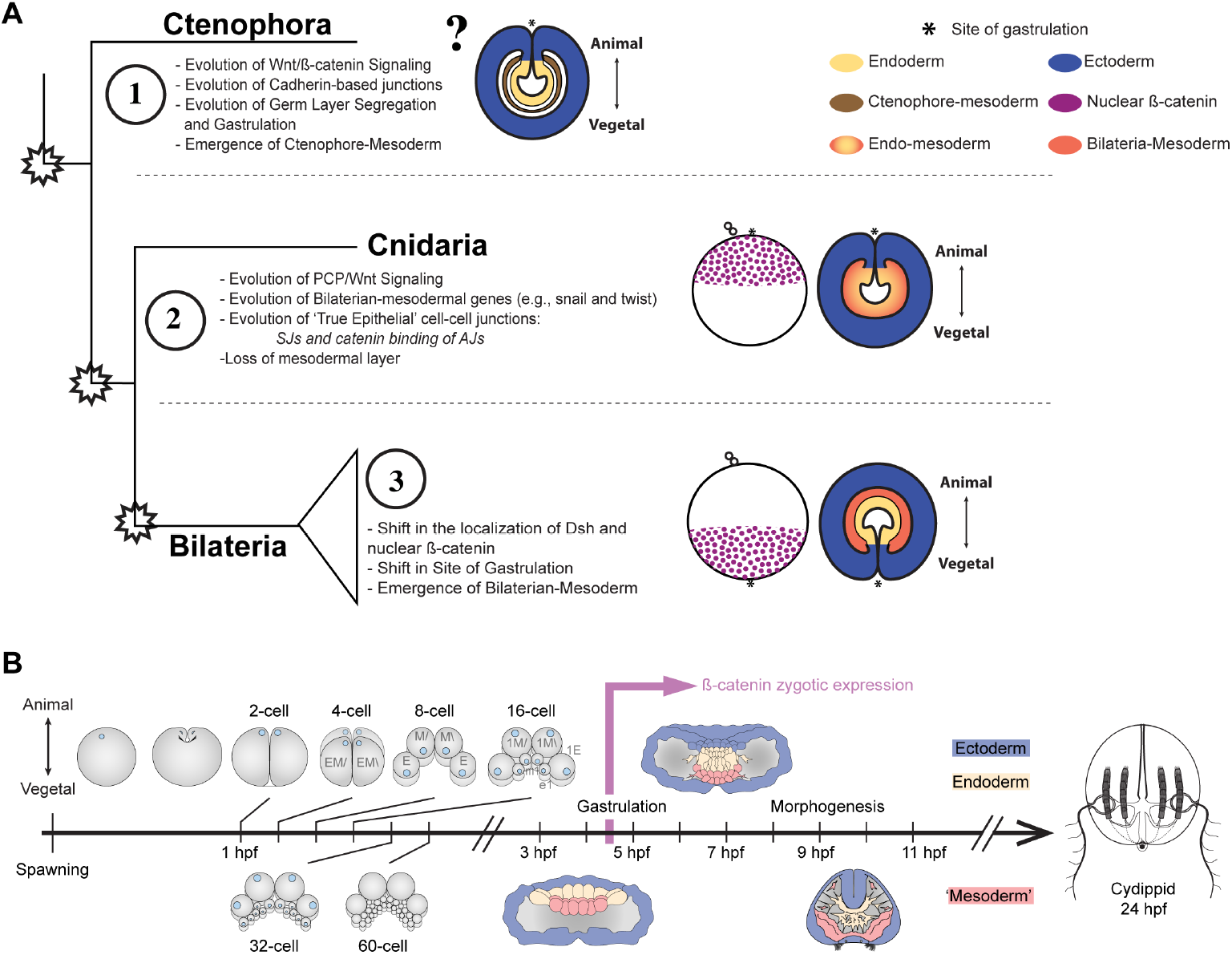
Evolution of Wnt signaling components during the animal evolution. A) Three major evolutionary steps (left side) that might have changed the organization of gastrulation in the Metazoa. The diagram (right side) depicts the site of gastrulation and the nuclear localization of ß-catenin during the development of Cnidaria and Bilateria. However, there are no descriptions available for ctenophore cells B) The stereotyped early development of *M. leidyi*. *M. leidyi* ß-catenin zygotic expression is temporally depicted.

Interestingly, the translocation of ß-catenin from the cytoplasm into the nucleus of blastomeres specifies the site of gastrulation in embryos of both bilaterian and cnidarian species (*4–17*). Thus, to understand how the subcellular localization of this protein is regulated has become a major topic of study in developmental biology research (*3, 8,18–26*). ß-catenin protein displays three different dominant cellular localizations: cytosolic, cortically attached to Adherens Junctions (AJs), and acting as a transcriptional co-factor inside of the nucleus (*19, 20*). Each of these localizations is regulated by specific molecular pathways whose components interact together as modules that organize cells in specific developmental contexts.

In most cells ß-catenin cytoplasmic levels are tightly regulated by its interaction with a destruction complex (GSK3ß/Axin/APC/CK1) where it is sequentially phosphorylated by CK1 and GSK3ß, which targets it for degradation in the proteasome (*19, 25*). On the other hand, when a Wnt ligand is present, cortical Dishevelled (Dvl) inhibits the action of GSK3ß and releases ß-catenin from its destruction complex. Free available cytosolic ß-catenin is either sequestered by apically polarized E-cadherin to the cell cortex or translocated to the nucleus. In the cell cortex, ß-catenin binds to the cytoplasmic domain of E-cadherin and alpha-catenin that forms the AJ belt complex required to connect these apical junctions to the cytoskeleton (*19, 25*). Nuclear ß-catenin has to interact with transcriptional factors such as LEF/TCF before inducing the expression of a variety of target genes. During bilaterian development cWnt signaling induces genes (e.g. snail, brachyury, twist) associated with the gene regulatory networks that determine endomesodermal cell fate (*8, 19, 25, 27*).

Recent work has indicated that the cWnt pathway was involved in endomesoderm specification in the last common ancestor to bilaterians and cnidarians (*19, 20*). Since ß-catenin also plays a critical role in cell adhesion, an outstanding question is if the ancestral role of this protein was in cell adhesion or in cell fate specification. Ctenophores provide an important model system to investigate this question. Ctenophores diverged prior to the cnidarian-bilaterian ancestor during animal evolution and the phylogenetic position of ctenophores may represent the earliest-branching extant animal clade (*28–31*). Ctenophores possess definitive muscle and mesenchymal cells in their mesoglea that are derived from endomesodermal precursors that cnidarians lack, thus, ctenophores provide a powerful opportunity to study the evolution of molecular mechanisms that may control ß-catenin localization and germ layer specification.

Despite the fact that genomic studies have shown the conservation of structural components of the ß-catenin protein in ctenophores (*32–34*), some upstream and downstream genes associated with its proper subcellular localization do not seem to be present in their genomes (*30, 33, 34*). For example, ctenophores do not appear to have the genes encoding for most cell-cell adhesion systems (*34, 35*), Wnt/planar cell polarity (PCP), and bilaterian-mesoderm (*30, 33*). Ctenophore cadherin does not have the cytoplasmic domains that bind to catenins (*30, 34*), and therefore, AJs cannot become functional in epithelial cells in the same way as it does in bilaterians and cnidarians. Moreover, ß-catenin zygotic expression starts after gastrulation is completed (*33*). Hence, the sequestration, degradation, and subcellular localization of ß-catenin could be quite different to what has been described for bilaterians and cnidarians.

All this together brings the question of what the ancestral role of ß-catenin might be in animal development and whether or not ß-catenin is involved in embryonic patterning as in other studied animals. Interestingly, a ß-catenin homolog has been described in the slime mold *Dictyostelium discoidin* (*36*) and has been implicated in the formation of the multicellular “slug” stage of their life cycle, suggesting that ß-catenin protein domains might have been involved in cell adhesion well before animals existed. Here, we developed a specific antibody (Figure S1; supplementary material) against *Mnemiopsis leidyi* ß-catenin (*Ml*ß-catenin) and analyzed its subcellular localization during the ctenophore *M. leidyi* embryogenesis. We have made fusion protein where *Ml*ß-catenin was fused to a fluorescent protein and expressed them in embryos by microinjection of mRNA so that we can visualize their localization in live animals. In addition, we tested the function of ß-catenin by the downregulation of its signaling. Regardless of the differences in the mechanism that regulate ß-catenin localization between ctenophores and (cnidaria+bilateria), we find no evidence that ß-catenin is involved in cell adhesion and our data suggest an ancestral role of nuclear*Ml* ß-catenin in tissue specification at the site of gastrulation during early *M. leidyi* gastrulation.

## RESULTS

### *M. leidyi* ß-catenin localizes in the nuclei of prospective endodermal cells

The ctenophore *M. leidyi* develops by a highly stereotyped and synchronous cleavage program (Figure 1B; detailed in supplementary material) (*37, 38*). The eggs of *M. leidyi* are polarized along the animal-vegetal axis and (*39, 40*) the first cleavage furrow is observable at the animal pole 45-60 minutes post fertilization at room temperature (25°C). Subsequent rounds of division occur every 10 to 20 min. Nuclear *Ml*ß-catenin protein (n*Ml*ß-catenin) is observable in the egg and in every blastomere until the 16-cell stage (Figures 2A, 2B, and S2). At this stage, n*Ml*ß-catenin is present in every cell except for the middle vegetal micromeres (‘m_1_ (*38*): prospective apical organ, tentacles epithelium, and nerve tissues) where *Ml*ß-catenin remains cytosolic (Figure 2A). Gastrulation in *M. leidyi* embryos takes place at the animal pole around 3 hpf. The presumptive ectodermal micromeres derived from the vegetal pole expand and migrate towards the animal pole covering the endomesodermal macromeres by epiboly. During these stages n*Ml*ß-catenin is present in the nuclei of animal macromeres that will develop into the endomesoderm (Figure S3A). While cells undergoing epiboly advance towards animal cells, cytosolic *Ml*ß-catenin becomes transiently nuclearized in some micromeres in closer contact with animal macromeres that will later become pharyngeal cell precursors (3 hpf; Figure S3A). Therefore, *Ml*ß-catenin nuclearization is regulated by maternally loaded components and may have cell-fate specification roles before the zygotic expression of this protein.

**Figure 2.**
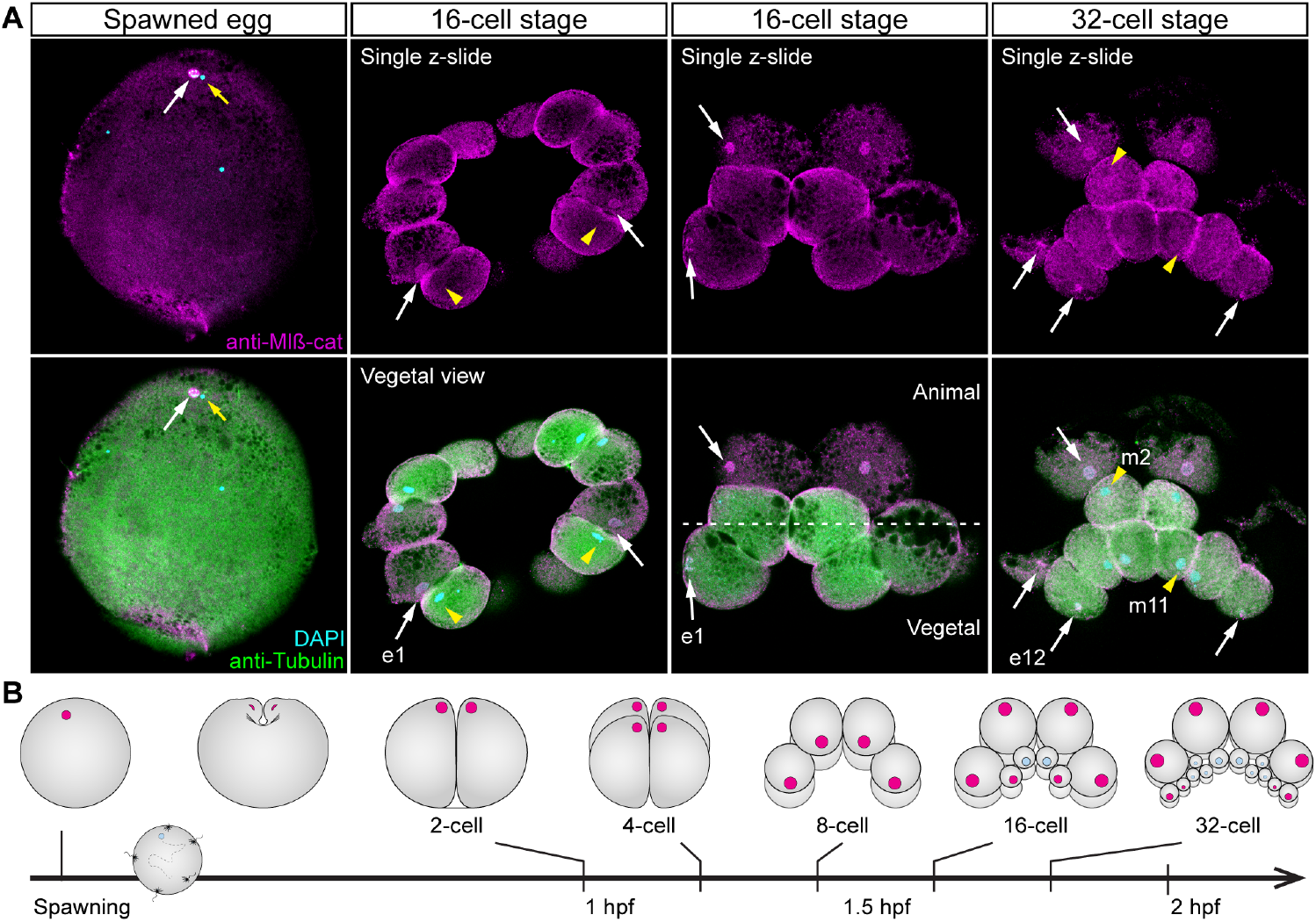
*Ml*ß-catenin protein localizes in the nucleus of the eggs and in every blastomere until the 16-cell stage. A) Immunostaining against *Ml*ß-catenin during cleavage stages of *M. leidyi* development. Nuclear *Ml*ß-catenin (white arrows) was not detected in the middle vegetal micromeres (yellow arrowheads) at 16-cell and 32-cell stages. Images are single optical sections from a z-stack confocal series. Orientation axes are depicted in the figure. B) Diagram depicting the nuclear localization of *Ml*ß-catenin. Animal pole is to the top as depicted in A. Morphology is shown by DAPI and Tubulin immunostainings. Homogeneous cytosolic staining was digitally reduced. See Figure S2 for further details.

During the formation of the prospective ‘mesoderm’ (4-4.5 hpf), *Ml*ß-catenin localizes to the nuclei of prospective endodermal cells but remains cytosolic in the ectoderm and prospective ‘mesoderm’ (Figure 3A and S3A). Interestingly, at the vegetal pole, n*Ml*ß-catenin is transiently present in a subset of cells parallel to the tentacular plane (4 hpf; Figure 3A). However, this vegetal nuclearization disappears by the end of the next round of cell division where n*Ml*ß-catenin is only observable in the endodermal cells located between the invaginated (prospective) ‘mesoderm’ and outer ectoderm (Figure 3A). Importantly, we never observed cortical localization of *Ml*ß-catenin at the level of cell-cell junctions during any of these stages. These observations together suggest that the Wnt/ß-catenin canonical pathway gets activated first at the animal pole and at the site of gastrulation regardless of the lack of zygotic Wnt and ß-catenin gene expression in those cells at those stages (as assayed by whole mount *in situ* hybridization) (*33*).

**Figure 3.**
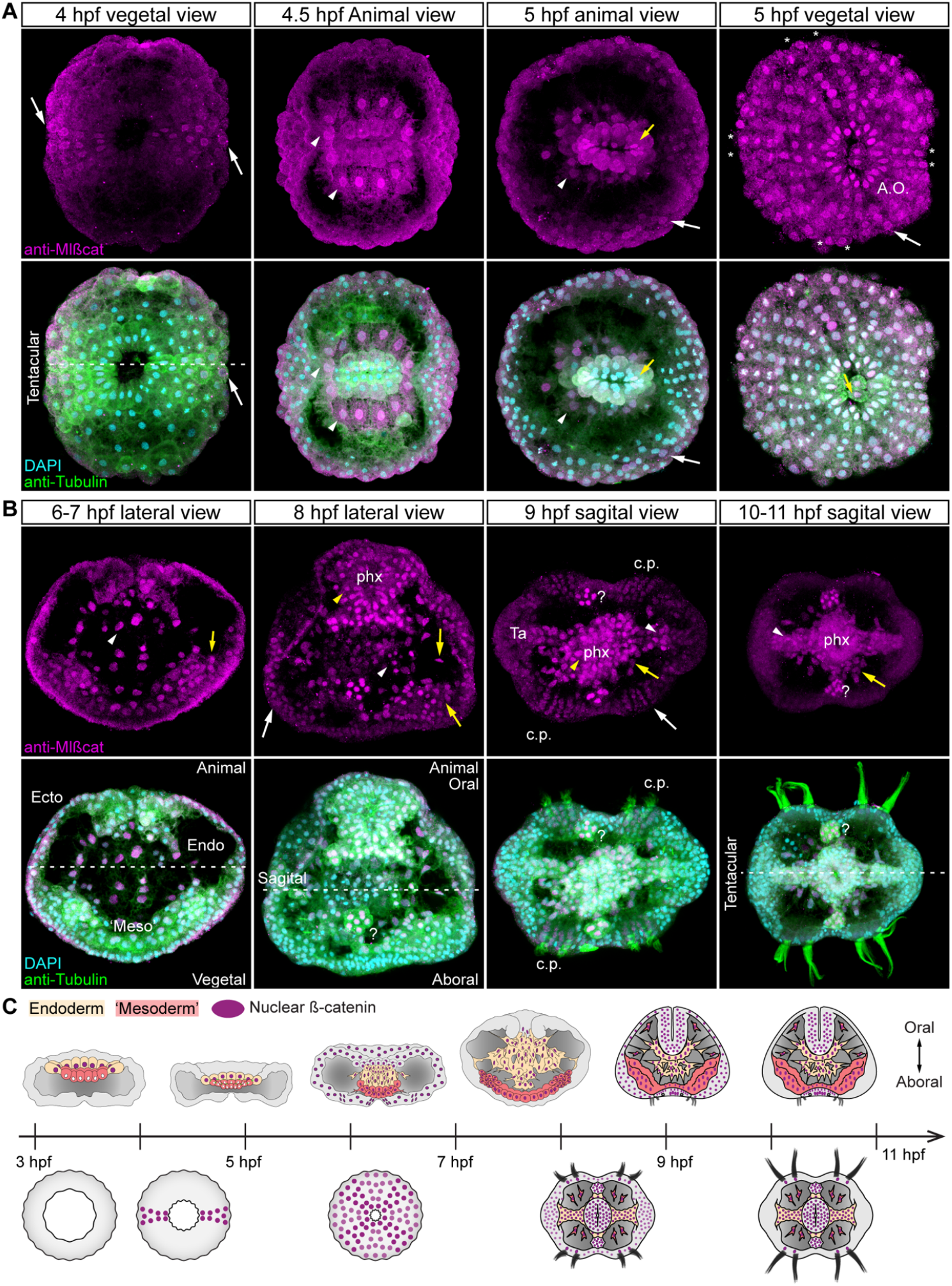
*M/ß*-catenin protein dynamics during and after *M. leidyi* gastrulation. A) Immunostaining against *Ml*ß-catenin during gastrulation stages of *M. leidyi* development. *Ml*ß-catenin protein localizes in the nuclei of prospective endodermal cells (white arrowheads). *Ml*ß-catenin protein is nuclear in subset of vegetal ectodermal cells at 4 hpf (White arrows). *Ml*ß-catenin protein transiently localizes in the nuclei of every cell at 5 hpf; white arrows indicate ectodermal cells. B) Between 6-7 hpf, nuclear *Ml*ß-catenin is only observable in endodermal (white arrowheads) and ‘mesodermal’ cells (yellow arrows). *Ml*ß-catenin protein transiently localizes in the nuclei of every cell at 8 hpf but it regionalizes after 9 hpf. *Ml*ß-catenin protein localizes in the nuclei of epidermal ectodermal cells (White arrows), pharyngeal cells (yellow arrowhead) endodermal cells (white arrowheads), and ‘mesodermal’ cells (yellow arrow). See Figure S3 for further details. Orientation axes and germ layers are depicted in the figure: Animal/Oral pole is to the top. Ecto: Ectoderm. Endo: Endoderm. ‘Meso’: ‘mesoderm.’ A.O.: prospective Apical Organ. ‘*’: prospective comb plates. phx: pharynx. Ts: Tentacle sheath. Ta: Tentacle apparatus. c.p.: comb plates. ‘?’: unknown tissue. No cortical localization was observed at the cell-contact region. Images are 3D reconstructions from a z-stack confocal series. Homogeneous cytosolic staining was digitally reduced. Morphology is shown by DAPI and Tubulin immunostainings. C) Diagram depicting the nuclear localization of *Ml*ß-catenin. Upper row: lateral view. Lower row: aboral view.

To confirm our observations, we cloned *Ml*ß-catenin (from 8-cell stage embryos) and developed an mRNA construct encoding for *Ml*ß-catenin fused to mScarlet (*Ml*ß-catenin::mScarlet) that was injected into zygotes, and the *in vivo* localization of the protein was recorded for the first time in *M. leidyi* embryos (Figure S4). The translated *Ml*ß-catenin::mScarlet was observed after 4 to 5 hours post injection, and therefore, it was impossible to observe this protein during early cleavage stages. During gastrulation, *Ml*ß-catenin::mScarlet was distributed uniformly in the cytosol and it was observed in the nuclei of oral macromeres (Figure S4) confirming our immunohistochemical observations (Figure 3 and S3).

### *M. leidyi* ß-catenin transiently localizes in the nuclei of every cell during and after gastrulation

At later stages of development, *Ml*ß-catenin remains cytosolic with no cortical localization observed (i.e. as would be expected if it were associated with cell-cell junctions) (Figure S3). During mid-gastrulation (5 hpf), when migrating ‘mesodermal’ cells are in direct contact with the ectodermal cells at the vegetal pole, n*Ml*ß-catenin is observable in all cells at apparently different concentrations (Figure 3A and Movie S1). At this stage, n*Ml*ß-catenin is highly concentrated in endodermal and ‘mesodermal’ cells, as well as, in some ectodermal cells: four pairs of cell rows in each embryonic quadrant and in a ‘ring’ of cells around the vegetal pole. Between 5-6 hpf, *Ml*ß-catenin remains cytosolic in all cells (Figure S3B) and ‘mesodermal’ cells start to migrate underneath the overlying vegetal ectoderm.

At 6 hpf (Figure 3B), n*Ml*ß-catenin is only observable in endodermal and ‘mesodermal’ cells until 8 hpf (Figure 3B). Intriguingly, between 8 and 9 hpf, n*Ml*ß-catenin is observable in all cells of the embryo at apparently different concentrations in ectodermal cells (Figure 3B). At this stage, n*Ml*ß-catenin is highly concentrated in endodermal and ‘mesodermal’ derivatives, as well as, in some ectodermal cells: pharynx, comb plates, and tentacle ectodermal-sheath. After 9 hpf, n*Ml*ß-catenin is only observable in all endoderm and mesodermally derived tissues and in the ectodermally derived pharynx, comb plates, and apical organ. In addition, we observed n*Ml*ß-catenin in two discrete regions positioned between the ctene-rows perpendicular to the tentacular axis (‘?’; Figure 3B), whose cell-lineage is still unknown.

### Downregulation of *Ml*ß-catenin signaling affects endomesodermal but not ectodermal differentiation

Previous studies have shown that *Ml*ß-catenin RNA transcripts are only detectable by *in situ* hybridization at the blastopore at 4 to 5 hpf (*33*), suggesting that zygotic expression starts after mesodermal cells have already internalized in the embryo. Later, *Ml*ß-catenin transcripts are ubiquitously expressed in the entire embryo (*33*). These observations suggest that *Ml*ß-catenin has maternal and zygotic roles during *M. leidyi* embryogenesis. Unfortunately, we do not have the technical capability yet to dissect the roles of maternal loaded proteins during the first stages of ctenophore development. Instead, we assessed the putative role of *Ml*ß-catenin after its zygotic expression by mRNA overexpression of recombinant Lv-cadherin (Figure 4) (*5, 12*) and CRRISPR/Cas9 knock-outs (Figure S7) delivered by microinjection into the uncleaved zygote.

**Figure 4.**
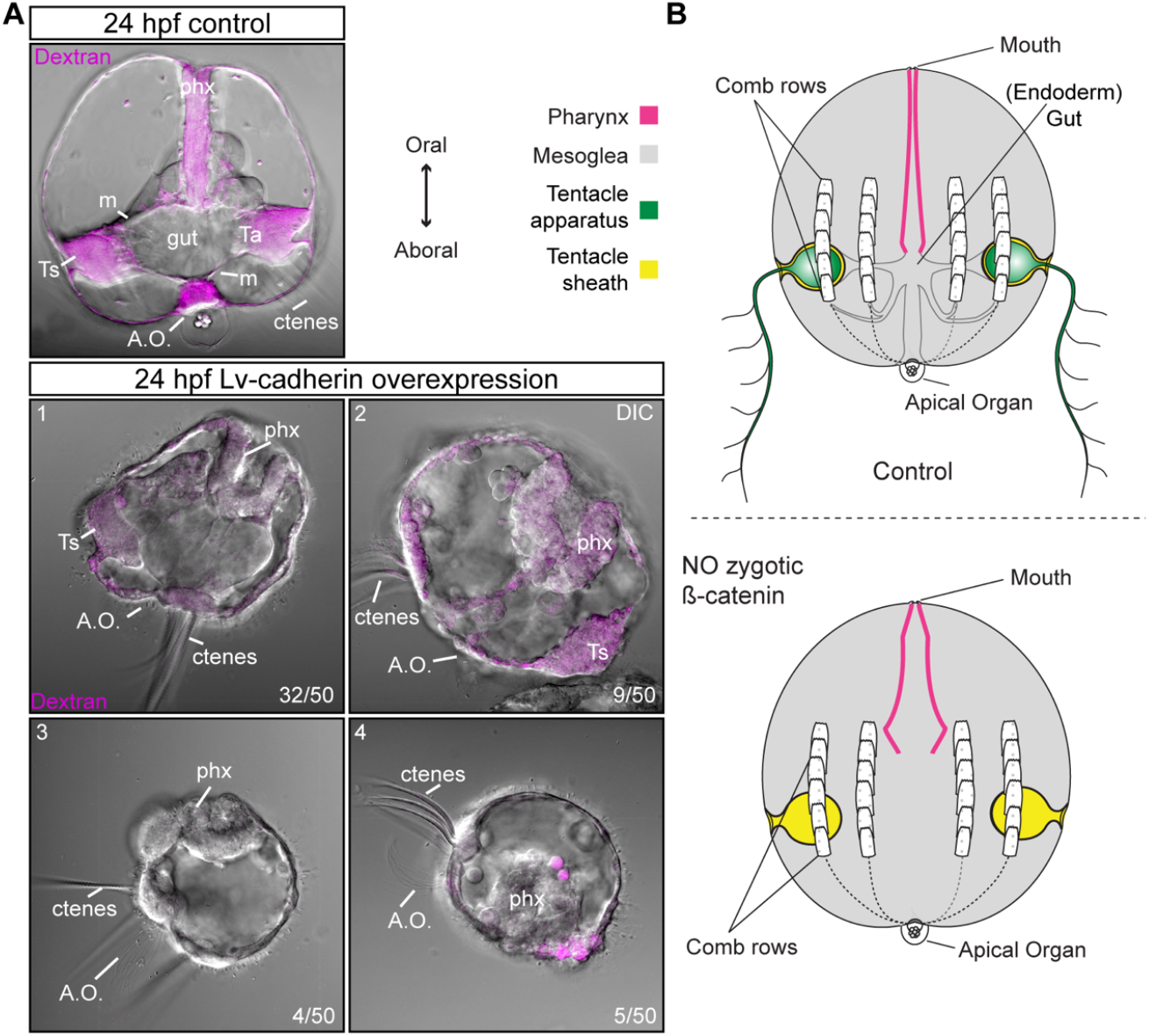
Testing the function of zygotic *Ml*ß-catenin. A) Overexpression of the mRNA encoding for the cytoplasmic domain of Lv-cadherin. Injected embryos lacked internal tissues including musculature (‘m’) and gut but developed ectodermal structures such as comb plates (ctenes), apical organ (A.O.), ectodermal pharynx (phx), and epidermal tissue as is depicted in B. Four slightly different phenotypes are shown in A. Ts: Tentacle sheath. Ta: tentacle apparatus. Similar results were obtained using CRRISPR/Cas9 knock-outs of **Ml*ß-catenin* gene (Figure S6).

The overexpression of this exogenous protein (cytoplasmic domain of Lv-cadherin) has been deployed in other systems to sequester cytoplasmic ß-catenin and inhibit its function (*5, 9, 12, 16, 41*). We did not observed changes in the cleavage pattern or phenotypes during earlier stages compared to controls (Dextran or Histone::RFP (Movie S2) injected embryos). Hence, embryos overexpressing Lv-cadherin where scored after 24 hpf at cydippid stage by DIC confocal *in vivo* imaging (Figure 4A). All of the Lv-cadherin injected embryos (50/50) lacked internal endodermal and ‘mesodermally’ derived tissues (gut and musculature) but developed ectodermal structures (comb plates, apical organ/dome cilia, ectodermal pharynx, and epidermal tissue) and some mesenchymal cells (Figure 5). Interestingly, some of the injected embryos (9/50) displayed slightly different phenotypes: shorter pharynx elongation and absence of tentacular sheaths (Figure 4A: panels 3 and 4). This is probably because due to different levels of expression Lv-cadherin. We obtained similar preliminary phenotypes using CRISPR/Cas9 knock-outs of *Ml*ß-catenin gene (supplementary material; Figure S7).

**Figure 5.**
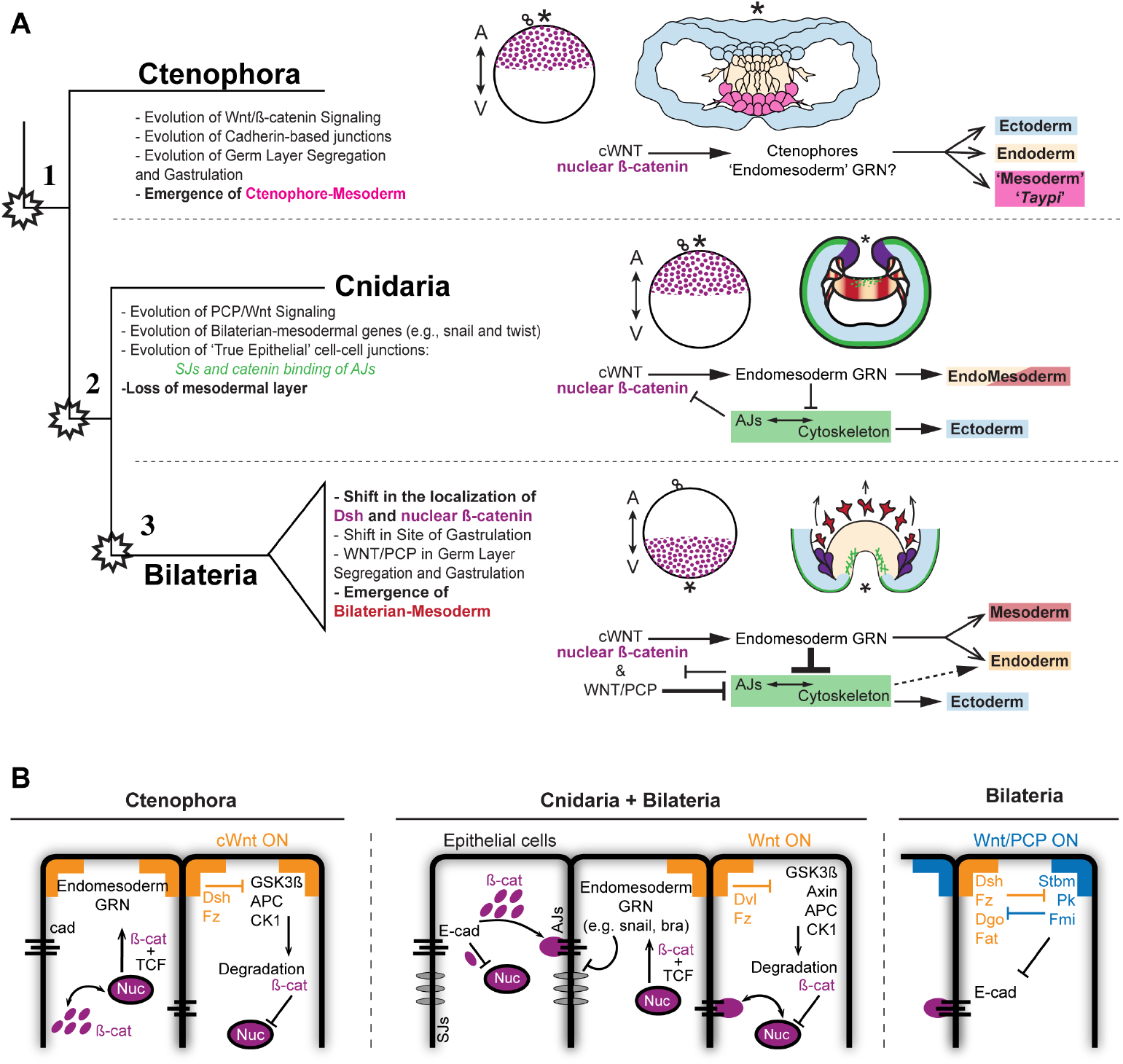
Evolution of ß-catenin regulation and gastrulation in Metazoa. A) The emergence of full sets of molecular modules rearranged the organization of cell-tissues that regulates the segregation of distinct germ layers. B) Diagram depicting the evolution of different interactions between known signaling pathways that ß-catenin localization in the Metazoa, including the new information obtained by this study during the early development of the ctenophore *M. leidyi.* Ctenophores do not have cell-cell adhesion complexes that regulate the localization of ß-catenin in cnidarian and bilaterian cells during gastrulation. In addition, the regulation of the Wnt/ß-catenin specification signaling by the Wnt/PCP signaling pathway appears to be present in bilaterian but not in cnidarians embryos.

## DISCUSSION

### The ancestral role of ß-catenin is cell-specification but not cell-cell adhesion

The structure and function of ß-catenin protein is highly conserved across all described metazoans. It was originally described, along with its closely related alpha catenin, to be associated with cell adhesion (*42, 43*) and only later shown to be involved in the nuclear activation of downstream target genes. The versatility of this protein (*19, 23, 25*) has now been thoroughly documented by its dual role as a component of AJs and as a transcription factor, in both adult and embryonic tissues, and thus has raised the interest of many researchers in different animal model organisms. In this sense, the recently sequenced genomes of the ctenophores *M. leidyi (*30**) and *Pleurobrachia pileus (*44**) have added another taxon to the long list of animals where ß-catenin primary structure is conserved. *Ml*ß-catenin possesses armadillo repeats flanked by the N-terminal (cortical and AJs binding) and C-terminal (nuclear role as transcriptional cofactor) characteristic domains (*32–34*).

Interestingly, the *M. leidyi* and *P. pileus* genomes do not contain the components for essential signaling pathways (e.g., Wnt/PCP, Notch, hedgehog) and bilaterian gene regulatory networks (e.g ‘mesodermal’ and neural GRNs) that regulate the nuclear signalization of ß-catenin protein in other metazoans (*30, 34, 35, 44*). Thus, due to the highly conserved function in bilaterians + cnidarians, the highly conserved protein sequence of ctenophore beta catenin, and the possible role of a ß-catenin-like armadillo gene involved in cell aggregation in slime molds (*36*), it strongly suggests the ancestral role of ß-catenin would be as a member of AJs and/or cell adhesion. However, recent studies have shown that ctenophore cadherin protein structure does not possess the catenin binding domains in its C-terminal domain (*34*). This prediction is borne out from our immunolocalization studies where we never see ß-catenin localized to cell junctions (Figures 2 and 3), as in the cnidarian *Nematostella vectensis* (*18, 22*) and other bilaterians (*5, 19, 26, 34, 45*). Hence, regardless of the conservation of *Ml*ß-catenin structure, *M. leidyi* tissues do not develop catenin-mediated AJs that mediate cell-cell adhesion.

Here, for the first time, we show the spatiotemporal localization of ß-catenin protein and the zygotic role of *Ml*ß-catenin protein during the early embryogenesis of the ctenophore *M. leidyi*. We show that *Ml*ß-catenin only displays cytosolic and nuclear localization during the observed developmental stages (Figures 2 and 3) and the zygotic protein is required for the differentiation of endo- and ‘mesodermally’ derived tissues (Figure 4). Therefore, we suggest that the ancestral role of ß-catenin is associated with cell specification and not with the formation of AJs or other roles in cell adhesion.

Moreover, the observation of n*Ml*ß-catenin and *Ml*Dvl (a cortical marker for Wnt signaling) in the zygote before and after the first cleavage at the animal pole (Figure 2 and S5) suggests that these proteins could be related to animal-vegetal axis specification displaying different maternal and zygotic roles (see extensive discussion in supplementary material). The localization of maternal n*Ml*ß-catenin in animal macromeres, and then, exclusively in endodermal cells (Figure 3; 4 hpf) suggest that maternal *Ml*ß-catenin has roles in specifying endomesoderm during early gastrulation (and hence, the site of gastrulation) between 3 to 4 hpf. These observations indicate that some of the developmental roles of maternally loaded ß-catenin and Dvl proteins are conserved across Metazoa (*4, 5, 14–17, 19, 25, 27, 6–13*).

### The interesting case of the ctenophore middle germ layer

More than the 95% of the extant animals are contained within the Bilateria. The ancestor of all bilaterian animals possessed three distinct germ layers: the ectoderm, endoderm, and mesoderm (*1, 2, 46*). Each germ layer is specified by a highly conserved set of GRNs and cellular mechanisms present across Bilateria and Cnidaria, even though the latter does not develop a distinct third layer (*8, 27, 47*). This fact brings the notion of tissues being organized by conserved molecular modules that interact together to organize cellular mechanisms orchestrating organismal development (*34, 48–50*). For example, in Bilateria and Cnidaria, the canonical Wnt/ß-catenin signaling pathway is finely regulated by the interaction between AJs and ‘mesodermal’ genes induced by the nuclear translocation of ß-catenin (*18, 19, 21, 25, 26, 45*).

Similarly, this study shows the first molecular evidence consistent with the presence of three different germ-cell layers characterized by the differential nuclearization of ß-catenin protein in a ctenophore embryo (Figure 3). Similar to bilaterian and cnidarian embryos (*4, 5, 14, 17, 51, 52, 6–13*), n*Ml*ß-catenin is present only in endodermal progenitor cells during early gastrulation (3-4 hpf) and can be used as a marker to identify the site of gastrulation and distinct germ layers. However, ctenophores do not have the AJs, signaling pathways ‘modules,’ nor the GRNs that contribute to mesodermal specification in their bilaterian and cnidarian relatives (*30, 34, 35, 50, 53*) bringing into question the homology of the middle germ layer in ctenophores. In other words, like bilaterians, ctenophores develop three distinct germ cell-layers, but they do it by different molecular mechanisms.

In this context, it is unclear how we establish homologies when we compare the specification of distinct germ layers across Metazoa. Is the differential localization of nß-catenin in the prospective endoderm during gastrulation definitive evidence that establishes germ layer homology? This would imply that, regardless the lack of genomic content in ctenophores, the role of nß-catenin specifying germ layers during early development is conserved across Metazoa. Thus, the cells of ctenophore germ layers could be considered to be homologous with other endomesodermal cells of bilaterians and cnidarians, but that the origin of mesoderm proper could be an independent event. The consideration of this option would also imply that cnidarians, whose genomes and cells possess the necessaries molecular and cellular mechanisms to segregate a distinct mesoderm (*18*), have lost the natural ability to develop a spatially distinct third germ layer.

If, on the other hand, we define the three distinct bilaterian-germ layers (ecto, endo, and mesoderm) on the basis of evolutionary conserved GRNs and or interactions between signaling pathways and not on morphological criteria, then, cnidarians but not ctenophores have germ layers homologous to bilaterians. This hypothesis would imply that the segregation of a distinct middle layer has two independent origins in Metazoa (*28, 30, 31*). In this scenario, it could have been possible that the emergence of full sets of molecular modules (e.g., AJs, SJs, Wnt/PCP signaling pathway, and mesodermal genes like twist and snail) rearranged the organization of cell-tissues (*30, 34, 35*). Such evolutionary event may have brought together new interactions between signaling pathways (e.g., junctional localization of ß-catenin) that would have led the emergence of a new epithelial tissue, the loss of a distinct third germ layer in cnidarians, and the later emergence of bilaterian-mesoderm (Figure 5).

In summary, we show that not only the structure but also the role of nß-catenin as a molecular marker to specify the site of gastrulation during early development is conserved across Metazoa (Figures 3 and S3). However, ctenophores do not have the mechanisms regulating the nuclearization of this protein that are necessary to differentiate distinct germ layers in other metazoans (*18, 30, 33–35*). Furthermore, the identity of the third distinct germ layer in ctenophore embryos might not be homologous to any other studied animal because of the absence of the GRNs that determine the bilaterian-mesoderm (*8, 27, 47*). Hence, until we identify the molecular components (and their functional interactions) that are deployed to tissue specification in ctenophores, we cannot discard any hypothesis with the available information. The development of temporal trajectories of single cell-seq and proteomic analyses of different cell lineages will give us the definite molecular candidates to experimentally test in a large phylogenetically scale. In the meantime, we suggest giving a new terminology to identify ctenophore ‘mesoderm’ (showed in quotation through the text) in order to highlight its different molecular identity. We suggest using the indigenous word *Taypi* (IPA: ‘tai.pi) that in the Aymara language (South American) refers to a central (middle) place of encounter (where two different entities encounter each other), reflecting the embryological nature of cells that reside in the “middle layer” in ctenophores.

## Acknowledgments

We thank C.E. Schnitzler and J. Ryan for technical assistance with the ctenophore genome.

## Author contributions

MS-S., and MQM. designed research and analyzed data with collaboration of AHW. MS-S. performed research with help of MQM. AHW design and developed *Ml*Dvl antibody. MS-S., and MQM. wrote the manuscript with collaboration of AHW. All authors read and approved the final manuscript.

## Data and materials availability

All data is available in the main text or the supplementary materials.

## SUPPLEMENTARY MATERIALS

### Materials and Methods

#### Culture and Spawning of *M. leidyi*

Spawning, gamete preparation, fertilization and embryo culturing of *M. leidyi* embryos was performed as previously described (*1*). Adult *M. leidyi* were maintained at the Whitney Laboratory for Marine Bioscience of the University of Florida (USA). Spawning was induced by incubating the adults under a three to four-hour dark cycle at room temperature. The embryos were kept in glass dishes (to prevent sticking) in filtered 1x seawater at room temperature until the desired stage.

#### Western Blot

Western blots were carried out as described (*2, 3*) using adult tissue lysates dissected by hand in order to discard larger amount of mesoglea. Antibody concentrations for Western blot were 1:1,000 for all antibodies tested.

#### Immunohistochemistry

All immunohistochemistry experiments were carried out using the previous protocol for *M. leidyi* (*1*). Embryos were fixed on a rocking platform at room temperature. Embryos of different stages were fixed for 2 hours in fresh Fixative (100mM HEPES pH 6.9; 0.05M EGTA; 5mM MgSO4; 200mM NaCl; 1x PBS; 3.7% Formaldehyde; 0.2% Glutaraldehyde; 0.2% Triton X-100; and 1X fresh sea water). Fixed embryos were rinsed at least five times in PBT (PBS buffer plus 0. 1% BSA and 0.2% Triton X-100) for a total period of 3 hours. PBT was replaced with 5% normal goat serum (NGS; diluted in PBT) and fixed embryos were blocked for 1 to 2 hours at room temperature with gentle rocking. Primary antibodies were diluted in 5% NGS to desired concentration. Blocking solution was removed and replaced with primary antibodies diluted in NGS. All antibodies incubations were conducted over night on a rocker at 4°C. After incubation of the primary antibodies, samples were washed at least five times with PBT for a total period of 3 hours. Secondary antibodies were then applied (1:250 in 5% NGS) and samples were left on a rocker overnight at 4°C. Samples were then washed with PBT and left on a rocker at room temperature for an hour. Samples were then washed once with PBT and incubated with DAPI (0.1μg/μl in PBT; Invitrogen, Inc. Cat. # D1306) for 1 hour to allow nuclear visualization. Stained samples were rinsed again in PBS two times and dehydrated quickly into isopropanol using the gradient 50%, 75%, 90%, and 100%, and then mounted in Murray’s mounting media (MMM; 1:2 benzyl benzoate:benzyl alcohol) for visualization. Note that MMM may wash DAPI out of your sample. We scored more than 1,000 embryos per each antibody staining and confocal imaged more than 50 embryos at each stage obtaining similar staining patterns for each case.

The primary antibodies and concentrations used were: mouse anti-alpha tubulin (1:500; Sigma– Aldrich, Inc. Cat.# T9026. RRID:AB_477593). Secondary antibodies are listed in Key resources table (Table 1).

Rabbit anti-*Ml*ß-cat antibody is custom made high affinity-purified peptide antibodies that were raised by Bethyl labs Inc. Affinity-purified *M. leidyi* anti-ß-catenin (anti-*Ml*ß-cat) peptide antibody was raised against a selected amino acid region of the *Ml*ß-catenin (METPVYQELS; Bethyl Inc., Montgomery, TX, USA). Rabbit anti-*Ml*Dvl antibody is custom made high affinity-purified against the DIX domain (recombinant protein) of *Ml*Dvl. Blast searches against the *M. leidyi* genome sequences showed that the amino acid sequences were not present in any predicted *M. leidyi* proteins other than the expected protein. Both antibodies are specific to *M. leidyi* proteins (Figure 2) and were diluted 1:100.

#### mRNA Microinjections

The coding region for *Ml*ß-catenin was PCR-amplified and cloned in frame with mScarlet into pCS2+ vector using NEBuilder^®^ HiFi DNA Assembly (NEB, Inc. #E2621S). Eggs were injected directly after fertilization as previously described for *N. vectensis* studies (*2–4*) with the mRNA encoding one or more proteins fused in frame with reporter fluorescent protein (N-terminal tag) using final concentrations of 300 ng/μl for each gene. Fluorescent dextran was also co-injected to visualize the embryos. Live embryos were kept at room temperature and visualized after the mRNA of the FP was translated into protein (4-5 hours). Live embryos were mounted in 1x sea water for visualization. Images were documented at different stages. We injected and recorded 20 embryos for each injected protein and confocal imaged each specimen at different stages for detailed analysis of phenotypes *in vivo.* We repeated each experiment at least five times obtaining similar results for each case. The fluorescent dextran and primers for the cloned genes are listed in Key resources table (Table 1).

#### CRISPR/Cas9 Knock-Outs

To target our gene of interest, we used synthetic guide RNAs (sgRNA; Synthego, Inc.) and followed the instructions obtained from the manufacturer to form the RNP complex with Cas9 (Cas9 plus sgRNAs). Target sites, off-target sites, and CFD scores were identified and sgRNA were designed using CRISPRscan (*5, 6*). We delivered the RNP complex by microinjection as previously described for *N. vectensis* studies (*3*). Lyophilized Cas9 (PNA Bio., Inc. Cat.# CP01) was reconstituted in nuclease-free water with 20% glycerol to a final concentration of 2μg/μl. Reconstituted Cas9 was aliquoted for single use and stored at −80°C. Embryos were injected, as described for mRNA microinjections, with a mixture (12.5μl) containing sgRNAs (80 ng/μl of each sgRNA), Cas9 (3 μg), and Alexa Fluor 488-dextran (0.2 μg/μl; Molecular Probes, Inc. Cat.# D22910). Cas9 and sgRNA guides only controls were injected alongside each round of experiments. sgRNA guides controls are only shown in figures as Cas9 had no significative effects when microinjected. 3 sgRNA were used to knock out *Ml*ß-catenin. We injected and recorded more than 20 embryos for each treatment. We repeated each experiment at five times obtaining similar results for each case. sgRNAs’ sequences and PCR primers flanking the targeted region are listed in Key resources table (Table S1).

### Supplementary Text

#### Normal Development of *M. leidyi*

The ctenophore *M. leidyi* develops by a highly stereotyped and synchronous cleavage program (Figure 1B) (*7, 8*). Many Sperm and eggs are subsequently released to the water column and fertilization takes place immediately at the time of spawning. The eggs of *M. leidyi* are polarized along the animal-vegetal axis and are fertilized by polyspermy (*9, 10*). Prior to syngamy, the egg pronucleus migrates in the cytosol, and after it ‘choses’ one of the spermatozoids, both pronuclei fuse and the first cleavage proceeds re-determining the animal-vegetal axis (*10*). The first cleavage furrow is observable at the animal pole 45-60 minutes post fertilization at room temperature (25°C), is unipolar, and holoblastic. Subsequent rounds of division are also unipolar and occur at the apical surface of each blastomere every 10 to 20 min. The second cleavage also passes through the animal-vegetal axis perpendicular to the first cleavage plane and gives rise to four symmetric quadrants (the EM blastomeres). The third cleavage is oblique and forms four end (E) blastomeres towards the vegetal pole and four middle (M) blastomeres at the animal pole. Each M blastomere undergoes two rounds of asymmetric division towards the vegetal pole, resulting in two small m micromeres (m1 and m2) and ‘leaving’ one M macromere at the animal pole. On the other hand, each vegetal E blastomere undergoes three asymmetric cell divisions, giving rise to three small e micromeres (e1, e2 and e3) at the vegetal pole and an E macromere displaced towards the animal pole. Cell divisions of micromeres continue, increasing the number of micromeres but maintaining the number of 8 macromeres at the animal pole. Gastrulation of *M. leidyi* embryos takes place about 3 hours post fertilization (hpf) by the migration towards the animal pole of e-descendant and m-micromere descendant micromeres that begin to envelop the macromeres via epiboly. The animal macromeres divide perpendicular to the oral-aboral axis forming a set of a subset of eight “oral” micromeres at the animal pole (prospective ‘mesoderm’) that remain attached to their mother macromeres (prospective endoderm) by their tubulin midbodies. At 4.5 hpf the eight “oral” micromeres born at the animal pole (prospective ‘mesoderm’) divide and migrate between endodermal precursors towards the vegetal pole making direct contact with the vegetal ectoderm at 5 hpf. Gastrulation in *M. leidyi* embryos ends when the prospective ectodermal pharynx starts to invaginate at the animal (oral) pole around 6 hpf. At this stage, the internal morphology of the embryo is drastically modified. The endodermal cells (oral macromeres) start to divide and lose their distinct cellular structure forming a syncytial layer. ‘Mesodermal’ cells acquire mesenchymal morphologies and migrate laterally along the vegetal pole underneath the vegetal ectoderm, in the space located between the syncytial endoderm and the ectoderm. At 7 hpf, cells continue dividing with ectodermal thickenings invaginating to give rise to the future tentacle bulbs. The comb plates form around at 8-10 hpf. During these stages, all juvenile structures are already developed and the cydippid stage hatches from the egg membrane at 18-20 hpf.

Another characteristic of ctenophore development is the high level of mosaicism (*8*). The site of the first cleavage is causally involved with the determination of the site of gastrulation and the future oral pole (*7, 9, 11*). Each round of cell division partitions the maternal determinants asymmetrically into specific blastomeres that will develop into specific tissues. In other words, the bauplan of ctenophores is a direct consequence of the cleavage program until the gastrulation movements reorganize the final structure of the embryo. For example, the four EM blastomeres correspond to the four quadrants of the larval and adult body plan. The four pairs of comb rows are largely derived from the four E blastomeres. The aboral micromeres give rise to the entire ectodermal-derived tissues (epidermis, apical organ, pharynx, nerve net, ctene plates, and tentacle sheath), the internal macromeres give rise to the endoderm (gut, endodermal canal system, anal canals, and photocytes), and the eight oral micromeres give rise the ‘mesoderm’ (the entire musculature and mesenchymal cells).

#### Antibody Specificity

Rabbit polyclonal affinity-purified antibodies against unique peptide regions of *M. leidyi* ß-catenin (Bethyl Laboratories, Inc.) and Dishevelled were used to determine their spatial and temporal expression at different developmental stages. Each antibody was characterized by Western blotting to establish its specificity (Figure S1). Western blots of *M. leidyi* adult extracts showed that the antibodies recognized different bands (Figure S1) for *Ml*ß-catenin (predicted size 99.5 KD; Figure S1A) and *Ml*Dvl (predicted size 68.9 KD; Figure S1A). Pre-adsorption of the *Ml*ß-catenin antibody with a tenfold molar excess of the antigenic peptide (used to generate and affinity purify the antibodies) resulted in the elimination of the appropriate-sized single band for *Ml*ß-catenin (Figure S1A). *Ml*Dvl antibody detected a single band around 100 KD (Figure S1A), suggesting a high phosphorylation state as in other systems (*12, 13*). In addition, whole-mount immunohistochemistry pre-adsorption experiments were performed to test the specificity of the *Ml*ß-catenin antibody by whole-mount immunohistochemistry. The staining pattern was strongly, but not completely, attenuated in early embryos when pre-incubated antibodies were used (Figure S1B).

#### *Ml*Dvl is differentially localized during *M. leidyi* embryogenesis and resembles the localization of n*Ml*ß-catenin during gastrulation

Dishevelled is a protein that is normally associated with Wnt receptors and is actively involved in ß-catenin activation. To observe the localization of *Ml*Dvl protein, whose gene expression is detectable from the beginning of development (*14*), we used a specific *Ml*Dvl antibody. *Ml*Dvl was observed to be homogenously distributed in the cortex of *M. leidyi* eggs as punctate aggregates (Figure S5). No clear asymmetric localization was detected until the formation of the first cell division where *Ml*Dvl localizes to the animal pole at the cortex surrounding the first and second cleavage furrows (Figure S5). This data suggest that fertilization may trigger the segregation of *Ml*Dvl to the future animal pole. Later, at the eight-cell stage, *Ml*Dvl localizes to the vegetal pole at the cortex surrounding the third cleavage furrows (Figure S5). However, no asymmetric localization of *Ml*Dvl was clearly detectable in the following stages up to gastrulation.

Similar to *Ml*ß-catenin, *Ml*Dvl is detectable through the cytoplasm of all embryonic cells between 3-4 hpf (Figure S6) and displays a nuclear localization (*15, 16*) in prospective endodermal cells. During the epiboly of the micromeres (3-4 hpf), *Ml*Dvl localizes to the cortex of the vegetal cells forming a ring-like structure (Figure S6). Between 4-5 hpf, *Ml*Dvl localizes to the cortex and nuclei of the ‘mesodermal’ cells in close proximity with the vegetal micromeres, and in the nuclei of endodermal cells (Figure S6). Furthermore, at this stage, *Ml*Dvl distributes with the dividing chromosomes of 4-8 rows of vegetal micromeres perpendicular to the tentacular plane (n*Ml*ß-catenin (Figure 3A) is present in cells between these rows), and with the dividing chromosomes of 8-16 animal micromeres parallel to the tentacular plane (Figure S6). This distribution pattern suggests a tissue specific regulation of *Ml*Dvl that may have roles in cell specification (*13, 15–17*).

From 5 to 7 hpf (Figure S6), *Ml*Dvl is nuclear at the endodermal and ‘mesodermal’ cells, as well as, in the animal (prospective pharynx) and vegetal ectoderm (prospective apical organ). Between 8-9 hpf, *Ml*Dvl is differentially localized in the *M. leidyi* embryo (Figure S6). *Ml*Dvl is cortically localized in ectodermal cells surrounding the prospective pharynx (oral) and in ectodermal cells of the developing apical organ and tentacular sheaths (aboral). While its specific subcellular localization is not clear in cells of the oral ectoderm, *Ml*Dvl asymmetrically localizes to the basal cortex of the cells at the aboral ectoderm (Figure S6). *Ml*Dvl is also asymmetrically localized to the basal cortex of endo- and mesodermally derived cells in contact with the overlying aboral ectoderm, suggesting an activation of the Wnt signaling pathway in these tissues (*14*). In addition, *Ml*Dvl displays nuclear localization in cells of the prospective pharynx, the prospective apical organ, and internal tissues derived from endo- and ‘mesodermal’ cells. After 10-hpf, the cortical localization of *Ml*Dvl is no longer observable and its nuclear localization is barely detected in few tissues: aboral end of the pharynx, tentacles apparatus, and apical organ (Figure S6). Interestingly, *Ml*Dvl is not present in the comb plates cells (Figure S6).

#### CRRISPR/Cas9 KOs

We characterized the phenotype of CRRISPR/Cas9 KOs embryos at the cydippid stage. We repeated the experiment five times, obtaining similar phenotypes with a similar penetrance each time (proportion: 8/20 embryos injected each time). ‘Mutant’ embryos lack all of their internal endodermal and ‘mesodermally’ derived tissues (Figure S7). On the other hand, ectodermally derived epidermal and tentacular sheaths, comb plates, pharynx, and apical organ (dome cilia) developed normally. This is surprising if we consider our previous observations where n*Ml*ß-catenin is present in cells of all three germ layers (Figures 3 and S3). At this stage, we could successfully obtain preliminary phenotypes using CRRISPR/Cas9 knock-outs of *Ml*ß-catenin gene. However, due to the difficulties presented in manipulating ctenophore eggs, we were not able to confirm these results in a large scale to genotype the mutations and further work and technology development is required.

### Supplementary Discussion

#### Maternal Wnt/ß-catenin signaling pathway and cell-fate specification

The regulation of the Wnt/ß-catenin signaling pathway plays a key role in cell-specification during the embryonic development of studied animals. Because we do not have specific antibodies to other Wnt related proteins, we do not yet know whether the proteins of these genes are maternally loaded into the egg, but our data suggest that the Wnt/ß-catenin pathway is activated during early stages by maternally loaded *Ml*ß-catenin and *Ml*Dvl proteins (Figures 2 and S5). Our immunostaining results shows that n*Ml*ß-catenin is localized in both oral macromeres and E blastomere-descendant micromeres but not in m-M blastomere descendant micromeres at the 16-cell stage (Figures 2 and S2) and during epiboly before gastrulation at 3 hpf (Figures S3A). Previous studies have shown that even though the normal development of *M. leidyi* is affected by drug-modifications of cell division patterns and protein translation during early stages, the cell-fate of some tissues (e.g., comb plates and apical organ) remains unaffected in drug-treated embryos (*8*). These observations, along with Lv-cadherin experiments and preliminary CRISPR/Cas9 phenotypes, suggest that n*Ml*ß-catenin has maternal roles specifying early cell-fates lineages.

Furthermore, transcriptomic data (*18*) and the asymmetric localization of *Ml*Dvl to the animal cortex and cleavage furrow of early zygotes (Figure S5) suggest that the Wnt/ß-catenin signaling gets activated around the time of fertilization. This is interesting because ctenophores fertilization is uniquely characterized by the migration of the egg pronucleus that ‘chooses’ a single sperm pronucleus out of many pronuclei scattered around the cortical cytoplasm (*10*). The migration of the egg pronucleus is mediated by astral microtubules network generated by the sperm (*19, 20*), and once both pronucleus fuse, the first cell division determine the animal-vegetal axis (*9, 10, 19*). It could be possible that sperm entry differentially localizes *Ml*Dvl and other proteins by cytoskeleton rearrangements in the egg cortex, and the egg pronucleus (containing n*Ml*ß-catenin) moves towards the future animal pole where *Ml*Dvl is localized at higher concentrations. However, further improvements in are required to elucidate this and other hypotheses about the role and interactions of maternal Wnt signaling pathway and the establishment of the animal-vegetal (oral-aboral) axis. We need better methods to observe the dynamic and function of these proteins during early stages of development. Ctenophore development is highly stereotyped, and cell-fates of some adult tissues (e.g., comb rows and apical organ) are already determined at the first cleavage cycles (*8, 21, 22*). It has been suggested that a “cleavage clock” regulates early cell-fate segregation and determination (*8*), in this context, it might be that the Wnt/ß-catenin signaling dynamics observed here are markers of this cell cycle timing. Therefore, we should not only study the molecular composition but also the cell behavior of blastomere cell-linages to understand the interactions underlying the regulation of *Ml*ß-catenin and cell-specification.

#### Zygotic Wnt/ß-catenin Signaling Pathway and Gastrulation Movements

Transcriptomic data suggest that all four *M. leidyi* Wnt genes are expressed during early stages (*18*). However, previous results show that these genes are only detectable by *in situ* hybridization in the vegetal pole after the completion of gastrulation (5-6 hpf) when ‘mesodermal’ and endodermal cells have already internalized (*14*). This suggest that there is a transition between maternal and zygotic Wnts transcripts that may influence their subsequent function. Furthermore, *in situ* hybridization results also show that, with the exception of MlFzdA and *Ml*Dvl, transcripts for most of the components of the Wnt signaling pathway express after (e.g., MlFzdB, MlSfrp, and MlTcf) or during (e.g., *Ml*ß-catenin) later stages of gastrulation (*14*). Thus, their zygotic proteins are not involved in early embryonic patterning but may have roles in later morphogenesis or cell specification events (e.g., endodermal canals, gut, and musculatures).

As with bilaterian embryos (*23–28*), *M. leidyi* Wnt proteins may act as morphogens that lead the gastrulation movements of ‘mesodermal’ cells towards the vegetal pole where *Ml*Dvl may bind with MlFzd at the cortex, allowing the nuclearization of *Ml*ß-catenin in proximal cells. The cortical localization of *Ml*Dvl at the vegetal pole during gastrulation (Figure S6), the cortical localization of *Ml*Dvl in ‘mesodermal’ cells adjacent to the vegetal Wnt expressing ectoderm (Figure S5; 4 hph), and the expansion of *Ml*ß-catenin nuclearization after ‘mesodermal’ cells contact the vegetal pole (Figure 3; 5 hpf) suggest that zygotic Wnt ligand proteins are present in vegetal cells at this stage of development (*14*). In addition, our experiments overexpressing Lv-cadherin (Figure 4), and preliminary CRISPR/Cas9 results (Figure S7), suggest a conserved role of zygotic Wnt/ß-catenin signaling pathway in mesendodermal precursors and that zygotic *Ml*ß-catenin has a conserved role in tissue specification in later stages.

Intriguingly, our results show transient waves of *Ml*ß-catenin nuclearization in all cells of the embryo (Figure S3). These observations suggest transient and stage-specific waves of either Wnt signaling pathway activation or inactivation. This is consistent with the redeployment of Wnt signaling pathways in other systems. For example, in *N. vectensis* there is an early nuclearization of ß-catenin that specifies endomesoderm (*12*, *29*), but at gastrulation Wnt signaling is activated at the blastopore to pattern the entire oral-aboral axis (*30–32*).

The discrete gene expression of Wnt ligands in regions associated with the aboral/vegetal pole, tentacle apparati, and apical organ (*14*), is concordant with n*Ml*ß-catenin (Figures 3, S3) and *Ml*Dvl (Figure S6) proteins localization in post-gastrulation stages. However, such gene expression patterns do not explain the activation of the Wnt signaling pathway when n*Ml*ß-catenin is observed in all cells of the embryo. One option is that the morphogen activity of Wnt ligands induces the activation of the signaling pathway in neighboring cells (*23–28*). In this scenario, the inactivation of this pathway should act as a regulator of n*Ml*ß-catenin (Figure S8), most likely, in a dose-dependent manner (*33, 34*). Unless unknown factors inhibit the *M. leidyi* Wnt/ß-catenin signaling pathway, the genomic absence of cadherin-catenin binding domains (*35*), Wnt/PCP components, Wnt antagonists (DKK, WIF, and CER), and the netrin related domain of MlSfrp (*14, 36*) suggest that this inhibition may happen downstream of *MlFzd.* However, it could also be possible that the activation of Wnt/ß-catenin signaling pathway is regulated by a differential expression of TCF and or GSK-3ß (*37–40*). We need to develop more specific antibodies and recombinant proteins to these key players in order to elucidate the role of the Wnt pathway in *M. leidyi* development.

At this point, we cannot discard the unlikely possibility that *Ml*ß-catenin has no functional role in *M. leidyi* development and its nuclear localization is a consequence of its conserved protein structure (*14, 35, 41*). Hence, we can now only say that *Ml*ß-catenin protein is differentially localized during *M. leidyi* development and can be used as a molecular marker to identify different rounds of cell-tissue specification, including the segregation of the three distinct germinal layers.

**Figure S1.**
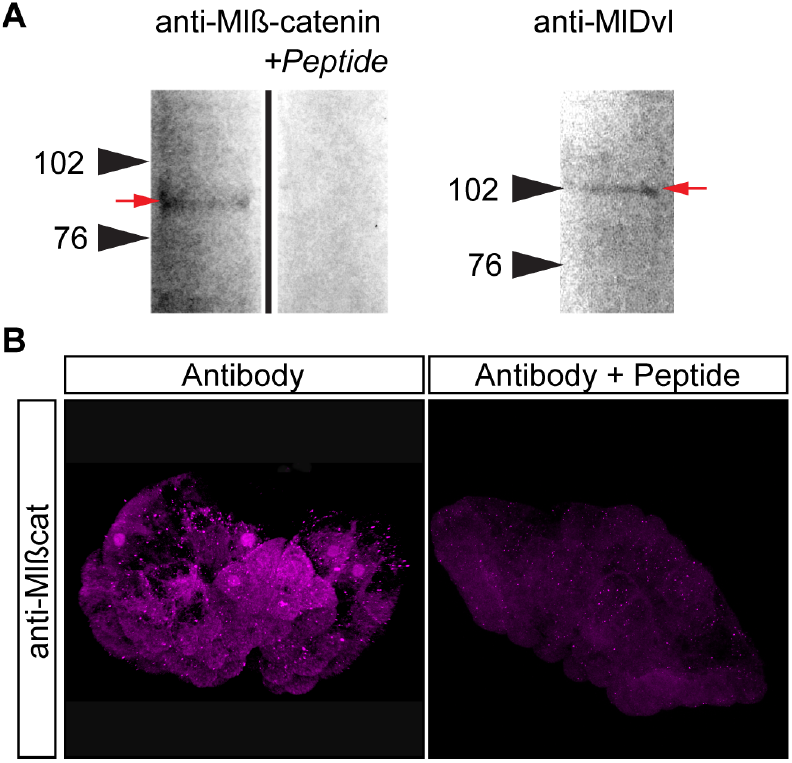
*M. leidyi* antibodies tested by western blots and pre-adsorption experiments. A) Western blots of *M. leidyi* adult tissue extracts using specific antibodies against *Ml*ß-catenin and *Ml*Dvl. Pre-adsorption of the antibodies with a tenfold excess of the antigen peptide resulted in the elimination of the staining of a single band for *Ml*ß-catenin. Arrowheads indicate the molecular weight in KD. A red arrow indicates the band recognized by the antibody for each protein. B) Whole-mount immunohistochemistry pre-adsorption experiments show that the staining pattern was strongly mitigated in early embryos when pre-incubated antibody against *Ml*ß-catenin with the respective peptide.

**Figure S2.**
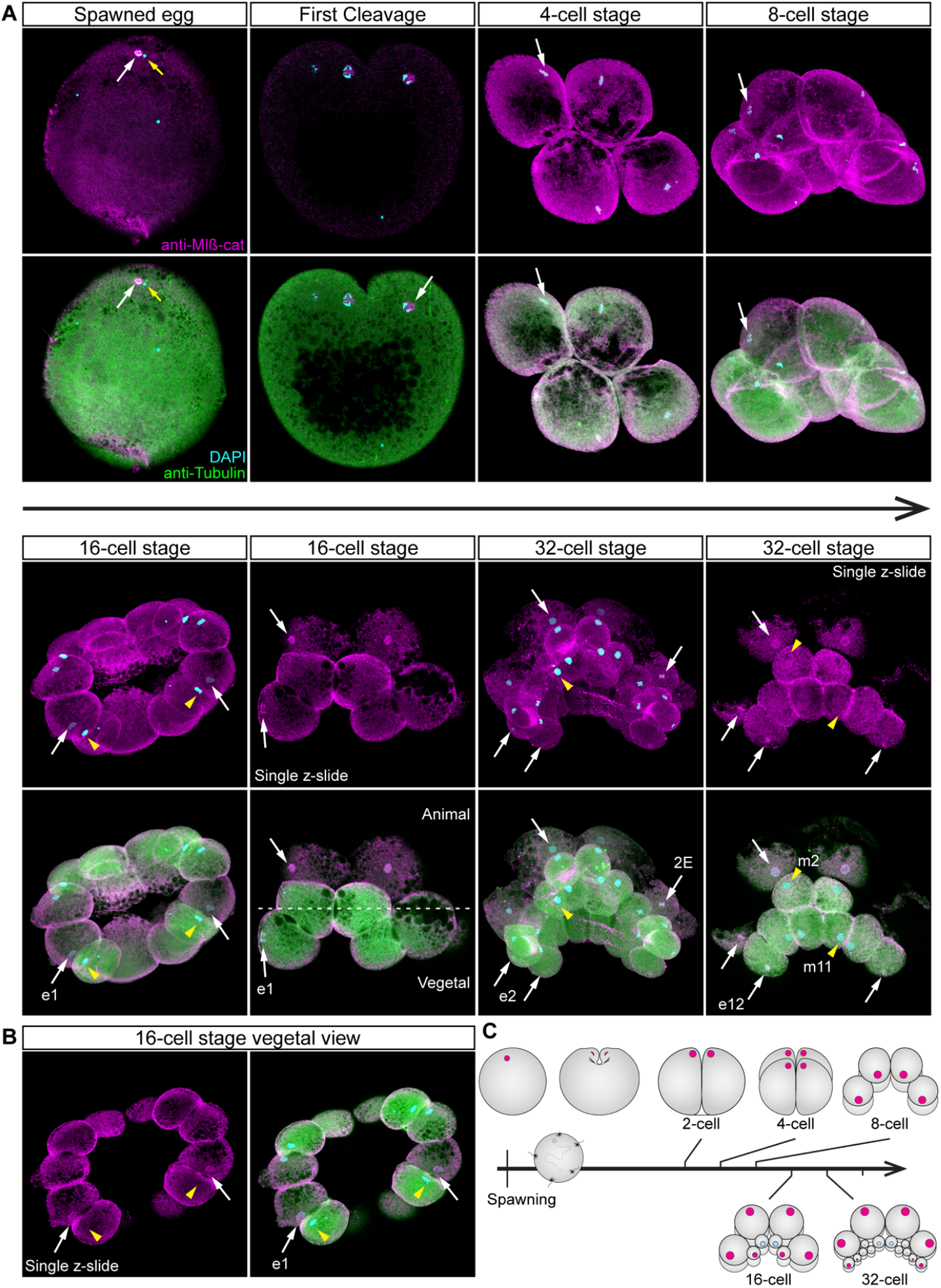
*Ml*ß-catenin protein localizes in the nucleus of the eggs and in every blastomere until the 16-cell stage. A) Immunostaining against *Ml*ß-catenin during cleavage stages of *M. leidyi* development. Nuclear *Ml*ß-catenin (white arrows) was not detected in the middle vegetal micromeres (yellow arrowheads) at 16-cell and 32-cell stages. Images are 3D reconstructions from a z-stack confocal series. Single optical sections are indicated. Orientation axes are depicted in the figure. B) A single optical section from the 16-cell stage z-stack confocal series shown in A. *Ml*ß-catenin protein localizes to the nucleus of e1 micromeres white arrows) but is absent from m1 macromeres nuclei (yellow arrowheads). C) Diagram depicting the nuclear localization of *Ml*ß-catenin. Animal pole is to the top as depicted in A. Morphology is shown by DAPI and Tubulin immunostainings. Homogeneous cytosolic staining was digitally reduced.

**Figure S3.**
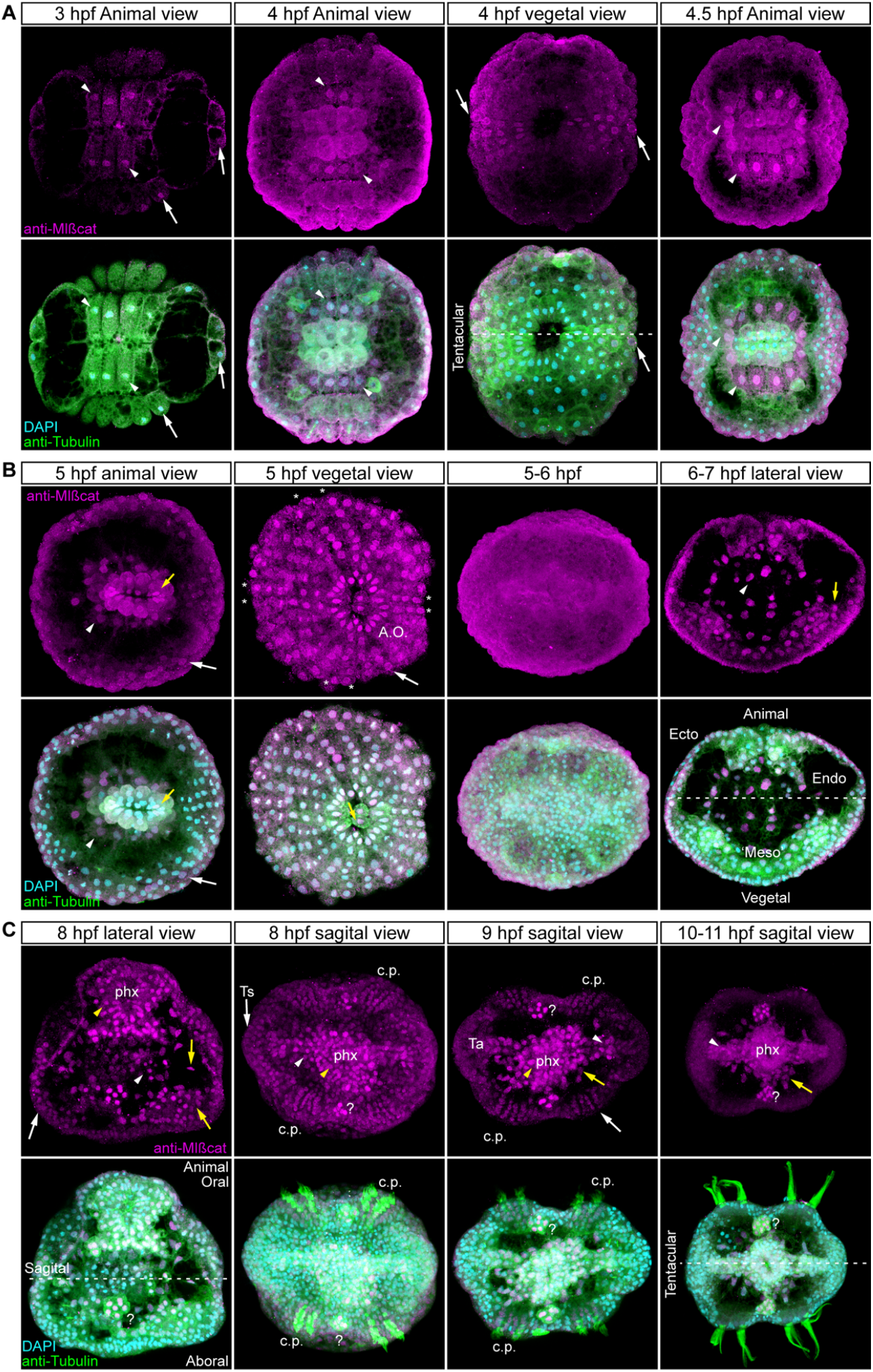
*Ml*ß-catenin protein dynamics during and after *M. leidyi* gastrulation. A) Immunostaining against *Ml*ß-catenin during early gastrulation stages of *M. leidyi* development. *Ml*ß-catenin protein localizes in the nuclei of prospective endodermal cells (white arrowheads). *Ml*ß-catenin protein is nuclear in the animal ectoderm at 3 hpf and in subset of vegetal ectodermal cells at 4 hpf (White arrows). B) Immunostaining against *Ml*ß-catenin during late gastrulation stages of *M. leidyi* development. *Ml*ß-catenin protein transiently localizes in the nuclei of every cell at 5 hpf; white arrows indicate ectodermal cells. Between 5-6 hpf, *Ml*ß-catenin remains cytosolic in all cells. Between 6-7 hpf, nuclear *Ml*ß-catenin is only observable in endodermal (white arrowheads) and ‘mesodermal’ cells (yellow arrows). Orientation axes and germ layers are depicted in the figure: Animal/Oral pole is to the top. Ecto: Ectoderm. Endo: Endoderm. ‘Meso’: ‘mesoderm.’ A.O.: prospective Apical Organ. ‘*’: prospective comb plates. C) Immunostaining against *Ml*ß-catenin after gastrulation stages of *M. leidyi* development. *Ml*ß-catenin protein transiently localizes in the nuclei of every cell at 8 hpf but it regionalizes after 9 hpf. *Ml*ß-catenin protein localizes in the nuclei of epidermal ectodermal cells (White arrows), pharyngeal cells (yellow arrowhead) endodermal cells (white arrowheads), and ‘mesodermal’ cells (yellow arrow). Orientation axes are depicted in the figure. phx: pharynx. Ts: Tentacle sheath. Ta: Tentacle apparatus. c.p.: comb plates. ‘?’: unknown tissue. No cortical localization was observed at the cell-contact region. Images are 3D reconstructions from a z-stack confocal series. Homogeneous cytosolic staining was digitally reduced. Morphology is shown by DAPI and Tubulin immunostainings.

**Figure S4.**
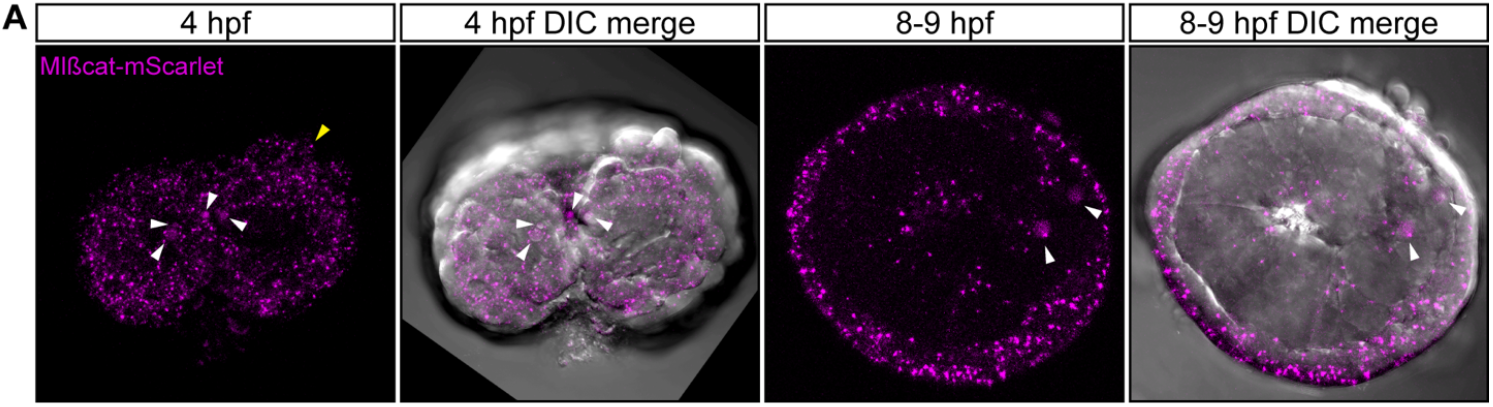
*In vivo* localization of *Ml*ß-catenin-mScarlet during different stages of *M. leidyi* development. The overexpression of *Ml*ß-catenin-mScarlet protein displays similar patterns observed with the antibody staining against the same protein at 4 hpf, and no cortical localization was observed. However, we were not able to dissociate nuclear *Ml*ß-catenin-mScarlet from its cytosolic localization by this method at later stages. All images are 3D reconstructions from a z-stack confocal series. Morphology is shown by DIC microscopy. White arrowheads indicate *Ml*ß-catenin-mScarlet protein nuclear localization. Yellow arrowhead indicates *Ml*ß-catenin-mScarlet protein cytosolic aggregates. Movie S3 shows the *in vivo* expression of Nvß-catenin::GFP into *M. leidyi* embryos that resembles antibody staining.

**Figure S5.**
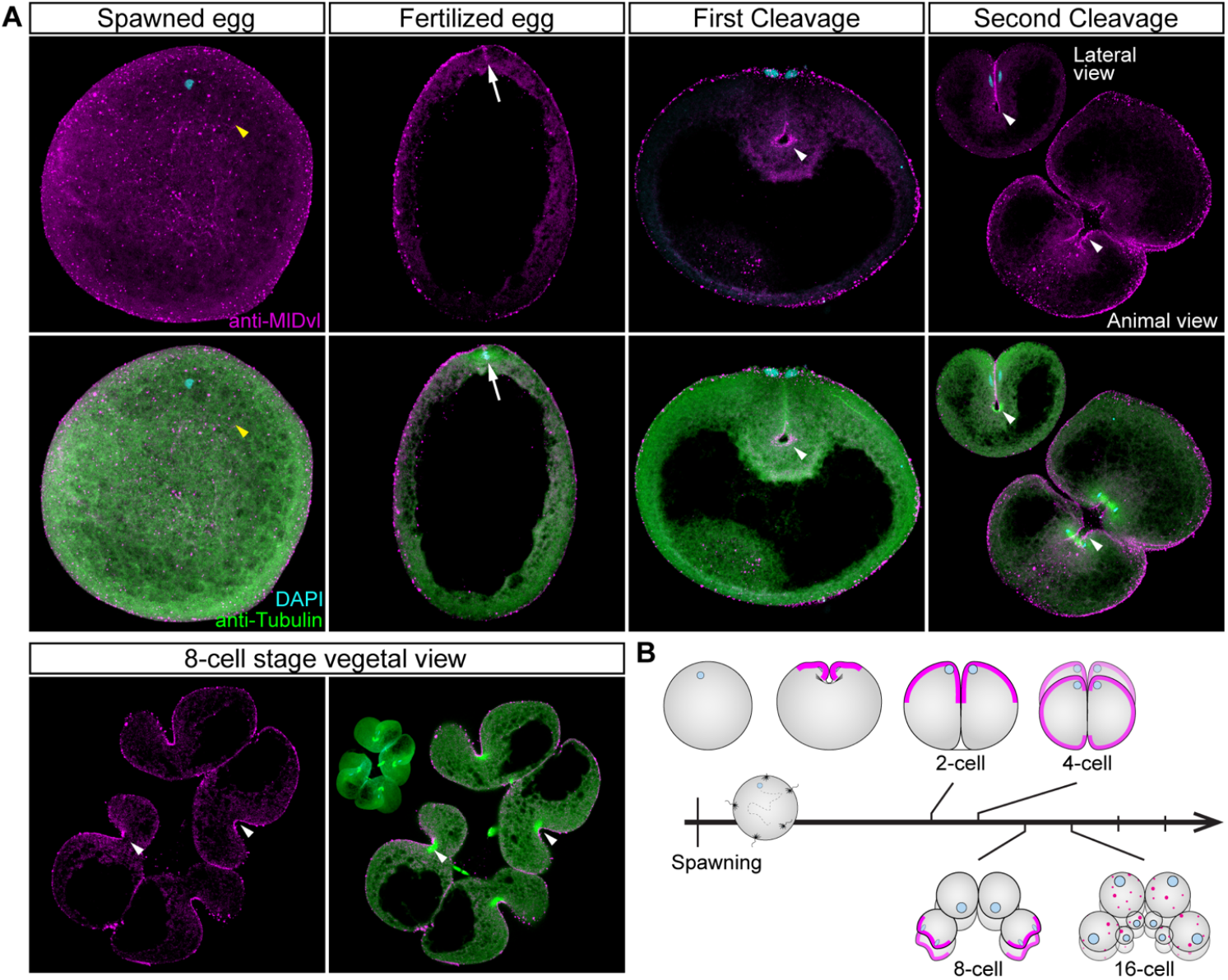
*Ml*Dvl protein localizes in the cortex surrounding the cleavage furrows. A) Immunostaining against *Ml*Dvl during cleavage stages of *M. leidyi* development. In the eggs, *Ml*Dvl protein distributes as punctate aggregates (yellow arrowhead). Cortical and asymmetric *Ml*Dvl protein (white arrow) was not detected until the first cell division. Orientation axes are depicted in the figure. Animal and lateral (upper left) views of two different embryos are shown in the second cleavage panels. Images are 3D reconstructions from a z-stack confocal series except for 8 cell-stage. 8-cell stage images are a single optical section from a z-stack confocal series (3D reconstruction is shown in the upper left corner for orientation purposes). White arrowheads indicate cortical *Ml*Dvl at the cleavage furrows B) Diagram depicting the cortical localization of *Ml*Dvl (magenta). Animal pole is to the top. Morphology is shown by DAPI and Tubulin immunostainings. Homogeneous cytosolic staining was digitally reduced. See Figure S5 for later stages.

**Figure S6.**
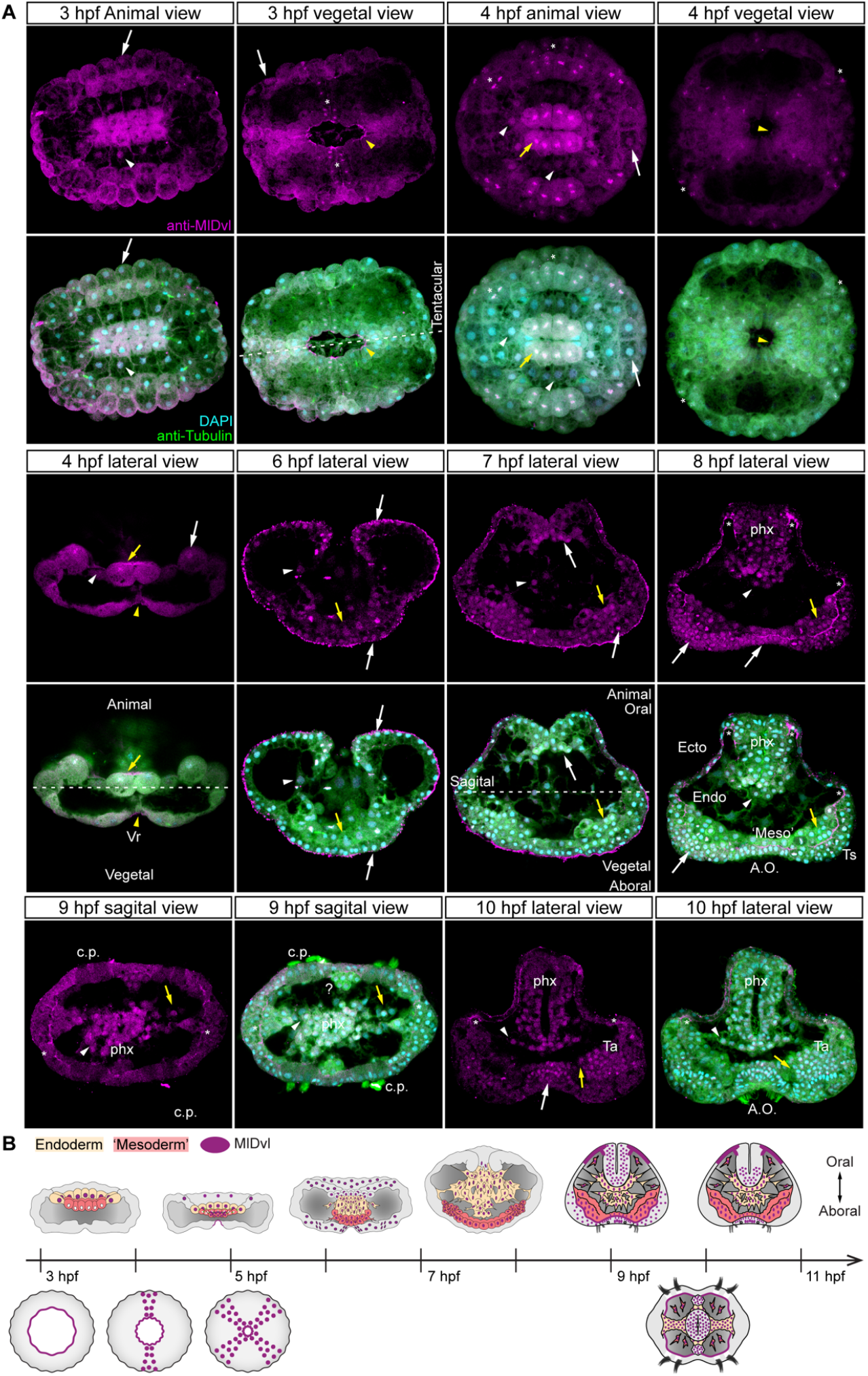
*Ml*Dvl resembles the localization of n*Ml*ß-catenin during gastrulation. A) Immunostaining against *Ml*Dvl during and after gastrulation stages of *M. leidyi* development. *Ml*Dvl protein localizes in the nuclei of prospective endodermal cells (white arrowheads) and ectodermal cells (white arrows). *Ml*Dvl protein distributes with the dividing chromosomes (‘*’) in the animal ectoderm and in subset of vegetal ectodermal cells at 4 hpf. *Ml*Dvl protein localizes to the cortex of the ‘vegetal-ring’ cells (yellow arrowheads). *Ml*Dvl localization in ‘mesodermal’ cells is indicated by yellow arrows. No cortical localization was observed at the cell-contact region. Images are 3D reconstructions from a z-stack confocal series. Orientation axes and germ layers are depicted in the figure. Ecto: Ectoderm. Endo: Endoderm. ‘Meso’: ‘mesoderm.’ phx: pharynx. Ts: Tentacle sheath. Ta: Tentacle apparatus. c.p.: comb plates. ‘?’: unknown tissue. ‘*’ at 3-4 hpf: *Ml*Dvl chromosomal localization. ‘*’ after 7 hpf: *Ml*Dvl cortical localization. B) Diagram depicting the *Ml*Dvl protein localization during and after gastrulation. Upper row: lateral view. Lower row: aboral view. Morphology is shown by DAPI and Tubulin immunostainings.

**Figure S7.**
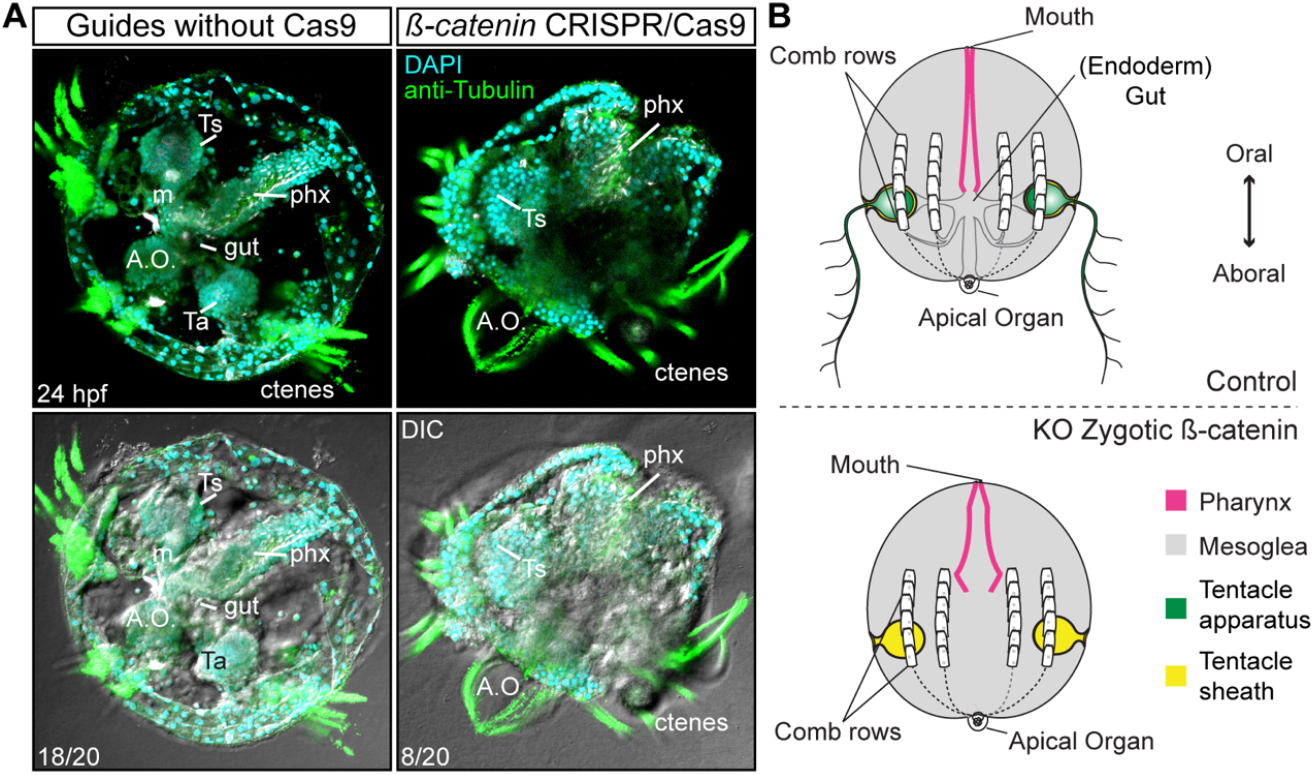
CRRISPR/Cas9 knock-outs of **Ml*ß-catenin* gene. A) Mutant embryos lacked internal tissues including musculature (‘m’) and gut but developed ectodermal structures such as comb plates (ctenes), apical organ (A.O.), ectodermal pharynx (phx), and epidermal tissue as is depicted in B. Ts: Tentacle sheath. Morphology is shown by DAPI and Tubulin immunostainings.

**Figure S8.**
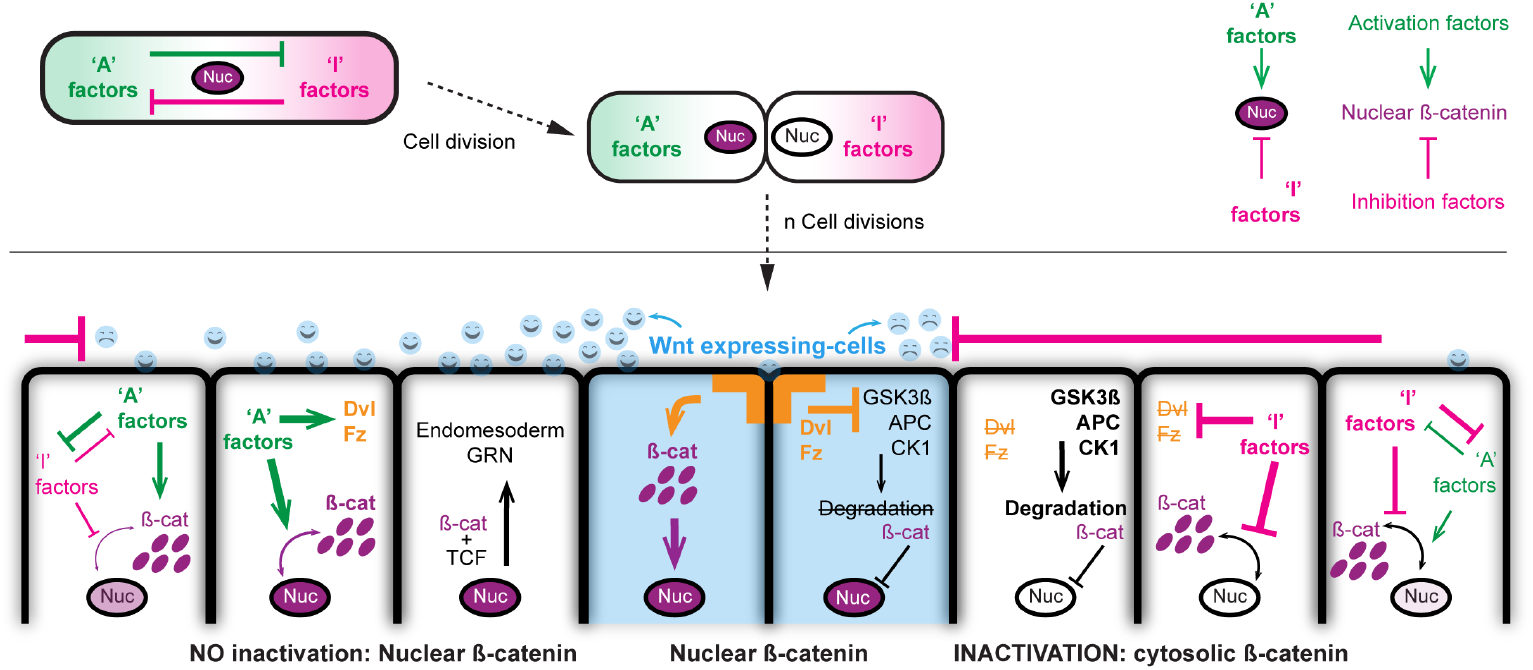
The morphogen activity of Wnt ligands during the early development of *M. leidyi.* Diagram depicting our hypothesis on the regulation of Wnt signaling and nuclear ß-catenin by the inactivation of this pathway in neighboring cells. *M. leidyi* cell-fate determinants are already polarized in the egg and partitioned since the first cell division. Several rounds of cell divisions distribute molecular factors that activate and or inhibit the Wnt signaling, leading to a differential nuclearization of Mlß-catenin.

**Movie S1**. Z-stack series of Immunostaining against Mlß-catenin during late gastrulation stages of *M. leidyi* development. Mlß-catenin protein transiently localizes in the nuclei of every cell at 5 hpf.

**Movie S2**. Z-stack series of *in vivo* Histone::RFP microinjected into the uncleaved eggs. Histone::RFP protein localizes in the nuclei of every cell at 24 hpf without altering the normal development of *M. leidyi* embryos.

**Movie S3**. Z-stack series of *in vivo* Nvß-catenin::GFP microinjected into the uncleaved eggs. Nvß-catenin::GFP protein resembles Mlß-catenin antibody staining.

---

**Table S1.**
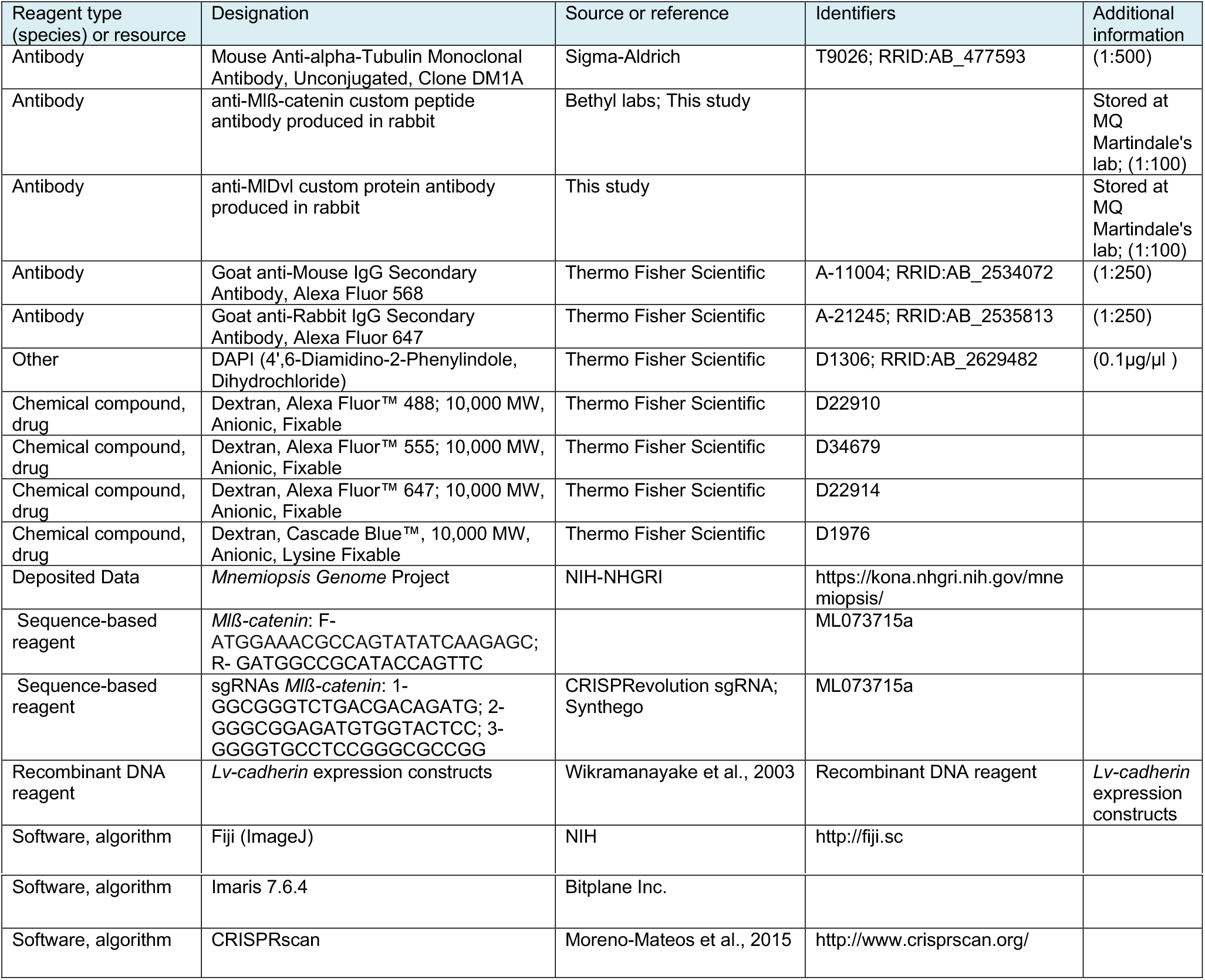
Key resources table.

