## Supplementary figures and images for "β-catenin has an ancestral role in cell fate specification but not cell adhesion"

### Movie S1

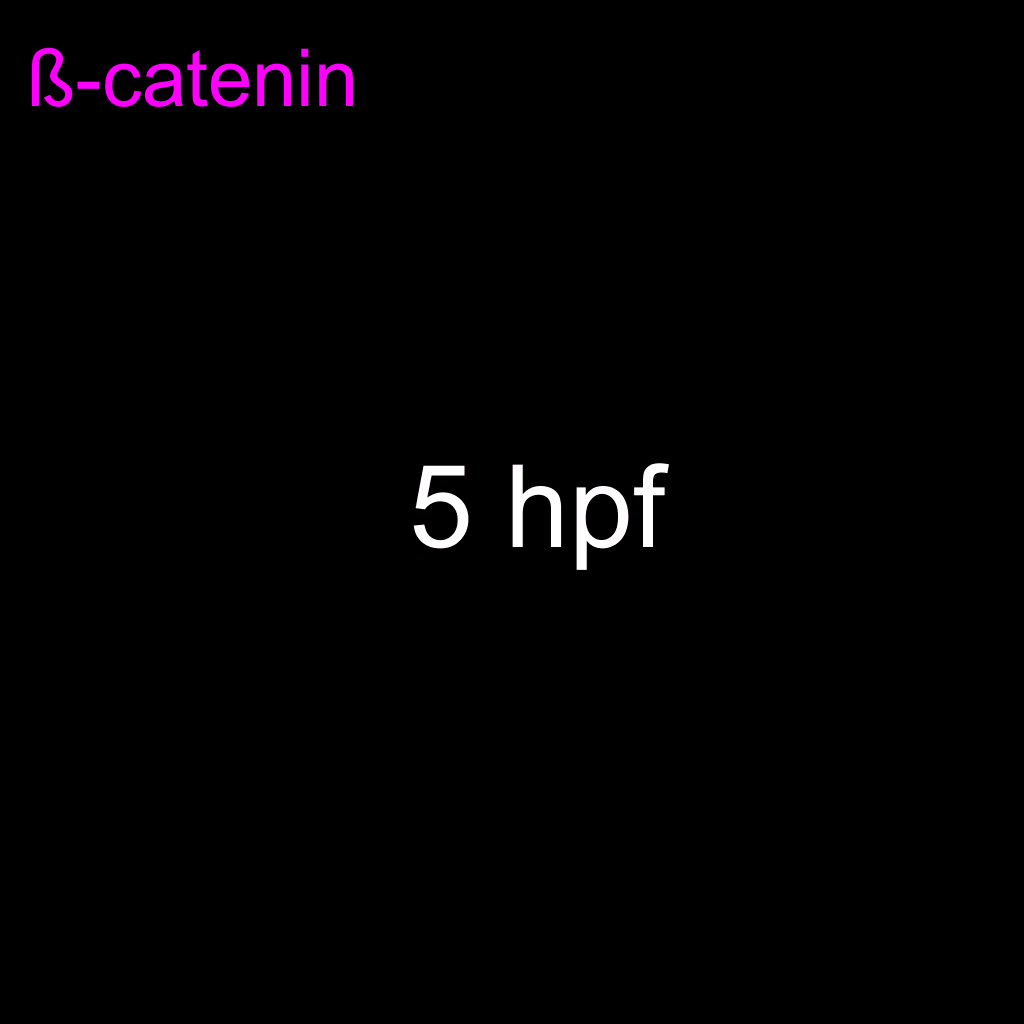

### Movie S3

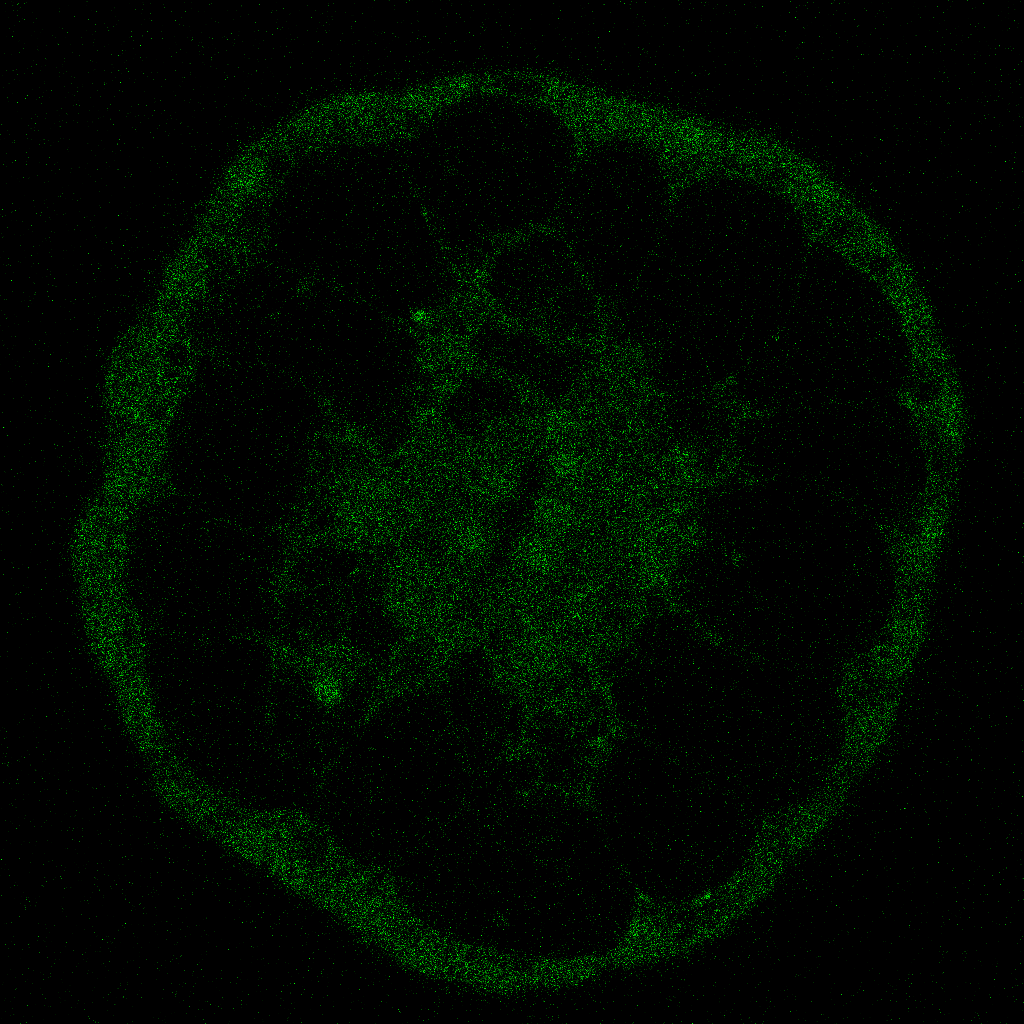
